# Structural basis for color tuning and passive ion conductance in red-shifted pump-fold channelrhodopsin ChR024

**DOI:** 10.1101/2025.09.16.675330

**Authors:** Yuka Takeno, Seiya Tajima, Suhyang Kim, Masaki Tsujimura, Yuma Ito, Masahiro Sugiura, Yo Yamashita, Koichiro E. Kishi, Masahiro Fukuda, Seiwa Nakamura, Masaya Watanabe, Hisako Ikeda, Kota Katayama, Yuji Furutani, Hideki Kandori, Hiroshi Ishikita, Keiichi Inoue, Hideaki E. Kato

## Abstract

ChR024 is a cation-conducting channelrhodopsin with a red-shifted absorption spectrum, recently discovered from a previously unidentified clade through machine-learning–guided gene mining. Due to its unique position in the phylogenetic tree and distinctive spectral properties, it serves as an important model for elucidating rhodopsin evolution and for engineering optogenetic tools with enhanced performance, but its structure and function remain poorly understood. Here, we present cryo-electron microscopy structures of ChR024 in detergent micelles and lipid nanodiscs at resolutions of 3.22 and 2.45 Å, respectively. The structures reveal an architecture strikingly similar to that of ion-pumping rhodopsins, even more so than other pump-like channelrhodopsins such as ChRmine. Structural, theoretical, and spectroscopic analyses uncover a unique color-tuning mechanism, in which charged residues close and distant from the retinal chromophore cooperatively modulate the pKa of the Schiff base counterion and thereby determine the absorption maximum wavelength. Finally, comparative structural analysis of channel- and pump-type rhodopsins, combined with electrophysiology, provides insights into the molecular boundary between these two functional classes and demonstrates the conversion of an outward proton pump into a light-gated channel. Together, these findings illuminate diverse mechanisms of color tuning and functional specification within the rhodopsin family, paving the way for the rational engineering of next-generation optogenetic tools.

## Introduction

Microorganisms have developed rhodopsin family proteins with diverse molecular functions and properties, allowing them to harness light energy to regulate various cellular activities. Among these, ion-translocating rhodopsins—microbial rhodopsins that function as ion pumps or ion channels—have been widely used as molecular tools since 2005–2006, enabling the light-dependent control of neuronal activity in living animals^1,2^. This technology, termed optogenetics, has revolutionized neuroscience and is now applied in broader contexts, including the regulation of osmotic pressure in plants, pH control in organelles such as mitochondria and lysosomes, and gene therapy for human eye diseases^3–5^.

To expand the scope of optogenetics, opsin genes have been isolated from microbial genomes and further optimized by protein engineering^6,7^. Mechanistic insights into ion-translocating rhodopsins have greatly facilitated both mining and engineering, driving the development of increasingly powerful optogenetics tools^8,9^. For example, inward H^+^ transport by rhodopsin was first achieved by introducing mutations into *Anabaena* sensory rhodopsin, a photochromic sensor^10^, and later confirmed by the discovery of a *bona fide* inward H^+^ pump, *Po*XeR, in nature^11^. Channelrhodopsins (ChRs) with markedly blue- and red-shifted absorption spectra were initially identified in natural sources^12–14^, but their *λ*_max_ values were subsequently extended through mutagenesis^15–17^. Similarly, anion-conducting channelrhodopsins (ACRs) were first engineered from cation-conducting channelrhodopsins (CCRs) using the C1C2 structure as a template^18–22^ and naturally occurring ACRs such as *Gt*ACR1 were later discovered^23,24^. A comparable trajectory was observed for potassium-selective channelrhodopsins (KCRs), which were first discovered in nature and subsequently refined through gene mining and protein engineering^25–28^. Despite these advances, our understanding of the molecular functions of ion-translocating rhodopsins and of the determinants that specify their diverse properties such as absorption spectrum, conductance, kinetics, sensitivity, and ion selectivity, remains incomplete.

Recently, a new ChR with a red-shifted absorption spectrum was identified through machine-learning (ML) –based genome mining and named ChR024^29^. This opsin belongs to a previously uncharacterized clade, phylogenetically distant from both canonical channel- and pump-type rhodopsins (Supplementary Fig. 1), yet functions as a robust CCR with one of the most red-shifted absorption maxima reported to date. Notably, because ChR024 occupies a phylogenetic position between known channel- and pump-type rhodopsins, and because of the unexpected gap between its experimentally determined absorption maximum (*λ*_max_ = 576 nm) and the ML prediction (*λ*_max_ = 543 nm) ^29^ , it provides a unique opportunity to investigate both the evolutionary relationship between channel- and pump-type rhodopsins and the mechanisms underlying red-shifted absorption in ChRs.

Here, we report high-resolution cryo-electron microscopy (cryo-EM) structures of ChR024 in both detergent micelles and lipid nanodiscs. Together with theoretical, spectroscopic, and electrophysiological analyses, these structures reveal the unique architecture and color-tuning mechanism of ChR024 and provide insights into the molecular principles that distinguish channel- and pump-type functions in ion-translocating rhodopsins.

## Results

### Structure determination of ChR024 in detergent micelle and lipid nanodiscs

To analyze the structural basis of color tuning and passive ion conductance, we first performed cryo-EM single-particle analysis of ChR024 (Methods, Supplementary Fig. 2). The full-length wild-type (WT) ChR024 (residues M1–T280) was expressed in Sf9 insect cells, purified in detergent micelles, vitrified, and imaged using a Titan Krios cryo-electron microscope (Supplementary Fig. 2a-c). The resulting structure in the ground state was determined at a nominal resolution of 3.22 Å (Supplementary Fig. 2d). The density map enabled the modeling of most ChR024 residues, but ambiguity remained in some side-chain rotamers and in densities potentially corresponding to water molecules. To improve resolution, we reconstituted purified ChR024 into MSP1E3D1 nanodiscs and repeated cryo-EM analysis, yielding a map at 2.45 Å resolution (Supplementary Fig. 2b-f). Although no major conformational differences were observed between the detergent- and nanodisc-embedded structures, the improved map allowed unambiguous modeling of amino acids, the retinal chromophore, associated lipids, and 16 water molecules (Supplementary Fig. 2g-k). Because no significant conformational differences were detected, the higher-resolution nanodisc-embedded structure was used for subsequent analyses.

### Overall structural comparison between ChR024, *Hs*BR, ChRmine, and C1C2

As in other microbial rhodopsins, ChR024 consists of a seven-transmembrane (7-TM) domain connected by three intracellular loops (ICLs) and three extracellular loops (ECLs), with a conserved lysine residue on TM7 covalently bound to the retinal chromophore (Fig. 1a). To characterize its structural features in detail, we compared ChR024 with three representative rhodopsins: bacteriorhodopsin from *Halobacterium salinarum* (*Hs*BR), a representative archaeal ion-pumping rhodopsin; ChRmine, one of the most potent optogenetics tools and a member of the recently identified pump-like channelrhodopsin (PLCR, also known as the bacteriorhodopsin-like channelrhodopsin, BCCR, family)^16,30^; and C1C2, a representative of the canonical channelrhodopsin family (Fig. 1b, c)^18^.

**Fig. 1.**
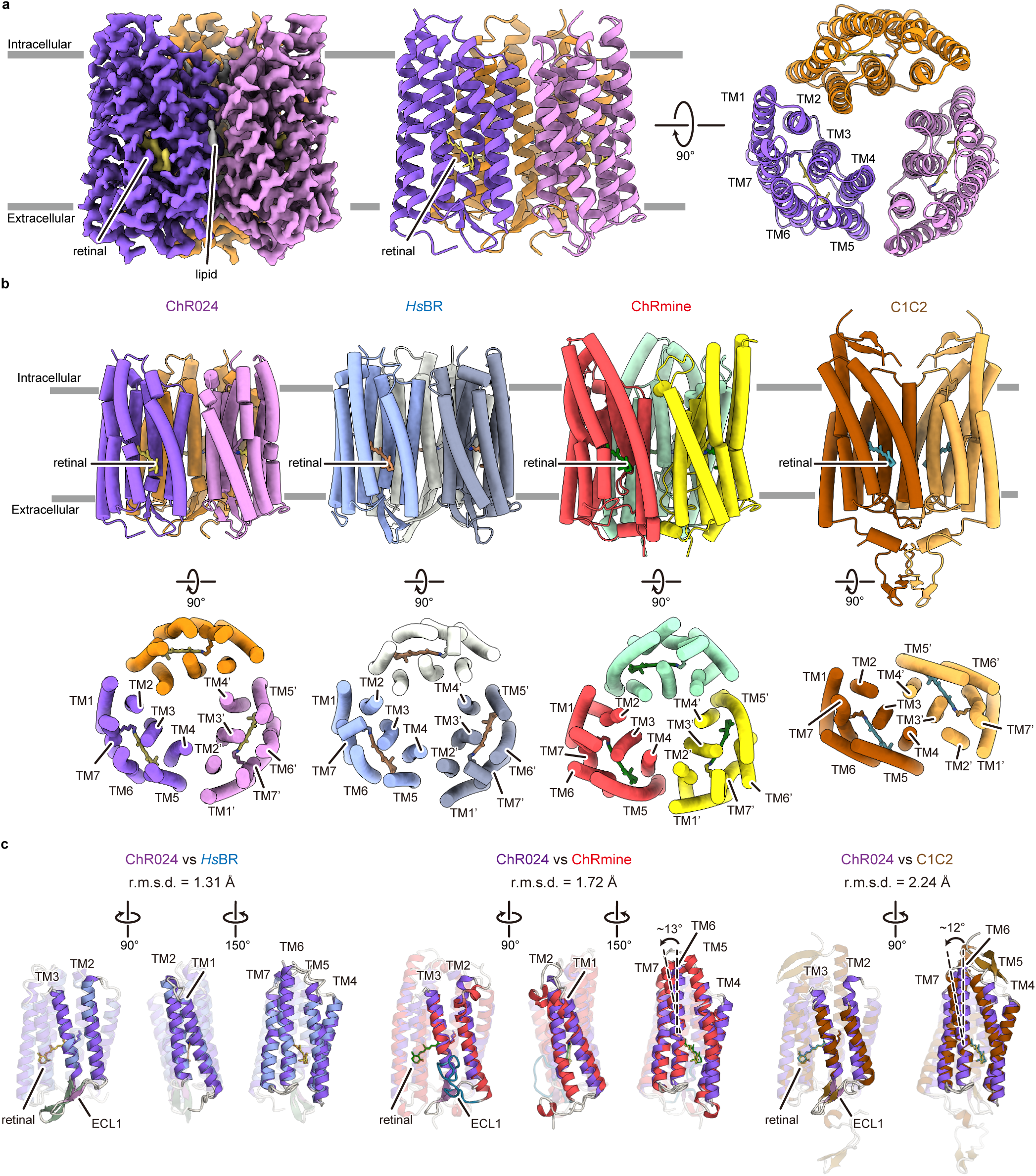
| Cryo-EM structure of ChR024 in nanodiscs. **a,** Cryo-EM density map (left) and ribbon representation of the ChR024 homotrimer viewed parallel to the membrane (middle) and from the intracellular side (right). Protomers are shown in purple, pink, and orange; the retinal chromophore in yellow; and lipids in gray. Dark gray bars indicate the approximate boundaries of the lipid bilayer. **b,** Overall structures of ChR024, a representative outward proton pump rhodopsin (*Hs*BR, PDB: 5ZIM),^42^ a representative pump-like/bacteriorhodopsin-like channelrhodopsin (PLCR/BCCR; ChRmine, PDB: 7W9W),^16^ and a representative canonical cation channelrhodopsin (C1C2, PDB: 3UG9)^18^ (from left to right). Structures are shown parallel to the membrane (top) and from the intracellular side (bottom). **c,** Structural comparisons of the ChR024 monomer with *Hs*BR (blue), ChRmine (red), and C1C2 (brown) from three different angles. ChR024 is colored according to the scheme in panel **(a)**. Retinals are shown in orange (ChR024), green (*Hs*BR), cyan (ChRmine), and brown (C1C2). ECL1 regions are highlighted in pink (ChR024), green (*Hs*BR), turquoise (ChRmine), and brown (C1C2). For clarity, certain transmembrane helices are shown in semitransparent colors depending on the viewing angle. Black dashed lines in panels c indicate the axes of TM6.

The first notable feature of ChR024 is that it forms a trimer, as observed in *Hs*BR and ChRmine, but in contrast to C1C2 (Fig. 1b). While all three proteins—ChR024, ChRmine, and *Hs*BR—form trimers, the assembly of ChR024 closely resembles that of *Hs*BR. In ChRmine, the relatively long N-terminus and tilted cytoplasmic end of TM2 create a large oligomeric interface (∼1,425 Å^2^) that stabilizes the trimer. By contrast, ChR024 and *Hs*BR have smaller interface areas (∼687 Å^2^ and ∼660 Å^2^, respectively), and lipid molecules intercalated between monomers likely play a more important role in maintaining their assemblies.

Beyond the oligomeric arrangement, the monomeric structure of ChR024 is also strikingly similar to *Hs*BR (Fig. 1c). All TM helices align closely with those of *Hs*BR, and even most ICLs and ECLs superimpose well. Notably, unlike ChRmine, ECL1 in ChR024 adopts a β-sheet conformation, again resembling that of *Hs*BR.

Taken together, although ChRmine belongs to the so-called “pump-like” or “bacteriorhodopsin-like” channelrhodopsin family, the tertiary and quaternary structures of ChR024 are in fact more similar to those of *Hs*BR.

### Retinal binding pocket and the Schiff base region

In rhodopsin proteins, the retinal chromophore is located at the center of the protein and is protonated in the dark. The positive charge of the protonated Schiff base is stabilized by one or two carboxylates on the extracellular side, historically referred to as counterions^31^. The absorption spectrum of rhodopsins is thought to be mainly influenced by three factors: (1) the planarity of the retinal chromophore, (2) the electrostatic potential around the retinal, and (3) the strength of the interaction between the Schiff base nitrogen and the counterion(s)^31–34^. To analyze the structural basis for the red-shifted absorption spectrum, we focused on the retinal binding pocket and the Schiff base region (Fig. 2a-c) and compared the ChR024 structure with those of *Hs*BR, ChRmine, and C1C2.

**Fig. 2.**
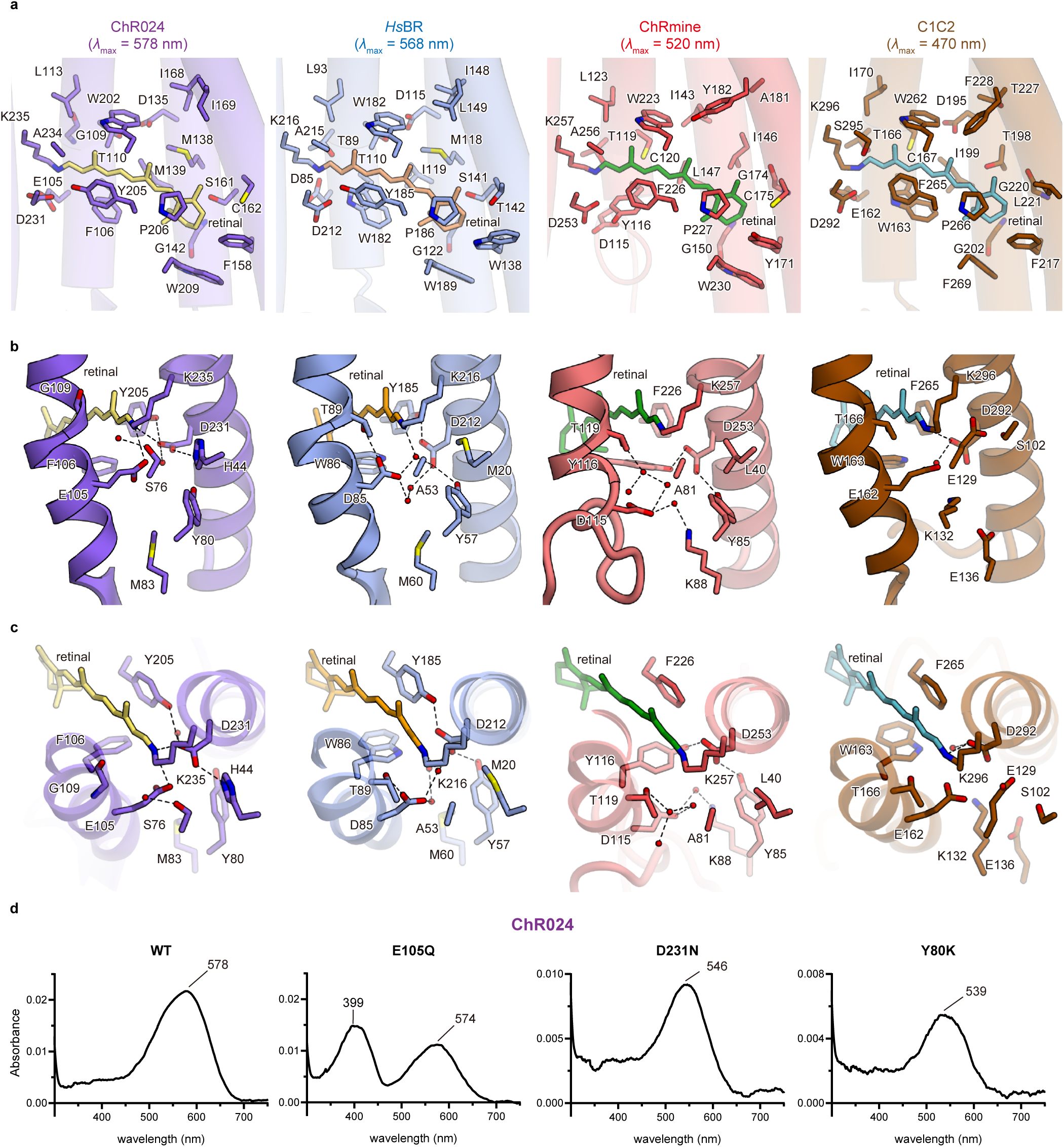
| Retinal binding pockets and Schiff base region. **a,** Retinal binding pockets of ChR024, *Hs*BR, ChRmine, and C1C2 (from left to right). Residues forming the retinal-binding pockets are shown as sticks. *λ*_max_ values represent the absorption maxima of each rhodopsin. Only glycine main chains are shown. TM5–7 are omitted for clarity. **b, c,** Comparison of the retinal Schiff base regions, viewed parallel to the membrane **(b)** and from the intracellular side **(c)**. Water molecules are shown as red spheres. Black dashed lines indicate hydrogen bonds with distances up to 3.2 Å. TM1, TM2, TM4, and TM6 are omitted for clarity. **d,** Absorption spectra of WT ChR024 and the E105Q, D231N, and Y80K mutants (from left to right) at pH 7.0.

First, we found that the overall retinal-binding pocket architecture is highly conserved, and the retinal maintains planarity within these pockets, although the 21 surrounding residues vary in sequence (Fig. 2a, Supplementary Fig. 3a). Notably, two residues near the β-ionone ring shift from less polar in C1C2 (G220/L221) to more polar in ChRmine (G174/C175), *Hs*BR (S141/T142) and ChR024 (S161/C162). By contrast, two residues near the Schiff base show the opposite trend, changing from polar in C1C2 (T166/S295) to non-polar in ChRmine (T119/A256), *Hs*BR (T89/A215), and ChR024 (G109/A234). Because polar residues near the β-ionone ring stabilize the excited S1 state and promote red-shifted absorption, whereas polar residues near the Schiff base stabilize the ground S0 state and promote blue-shifted absorption^31,33,35^, these sequence patterns are consistent with the progressively red-shifted *λ*_max_ values across the proteins. Another intriguing feature is the cysteine–aspartate pair on TM3 and TM4 (C167 and D195 in C1C2), commonly referred to as the DC gate. In several CCRs, including *Cr*ChR2 and C1C2, this pair is interconnected by water-mediated hydrogen bonds, and mutations at this site markedly extend the lifetime of the channel open state^36,37^. In ChR024, however, the cysteine is replaced by threonine (T110), which forms a direct hydrogen bond with D135 (Supplementary Fig. 3b). The T-to-C mutation does not accelerate channel kinetics^29^, and the interaction network more closely resembles that in *Hs*BR. This suggests that the function of the DC pair in ChR024 mutant is different from that in other ChRs and may be similar to that of *Hs*BR, implying a distinct gating mechanism for ChR024.

ChR024 also exhibits unique features in the Schiff base region (Fig. 2b, c). Unlike *Hs*BR or ChRmine, which contain two aspartates on TM3 and TM7 as Schiff base counterions, ChR024 has a glutamate (E105) on TM3, and the distance between E105 and D231 on TM7 is unusually short (2.7 Å) (Supplementary Fig. 3c). Furthermore, the residue one helical turn above the TM3 counterion, typically a serine or threonine (T89 in *Hs*BR and T166 in C1C2), is replaced by glycine (G109), creating extra space adjacent to the Schiff base nitrogen. This space is occupied by a water molecule coordinated by S76 (Fig. 2b, c). In addition, while the TM7 counterion commonly forms hydrogen bonds with one or two tyrosines on TM2 and/or TM6, thereby modulating its p*K*_a_, in ChR024 D231 does not form a hydrogen bond with Y80 but instead interacts with H44 on TM1 (Fig. 2b, c). Together, these features establish a hydrogen-bonding network in the Schiff base region that is strikingly different from those in *Hs*BR, ChRmine, and C1C2, which may explain why the proton is not efficiently released from the Schiff base nitrogen during the photocycle^29^.

The short distance between E105 and D231 is within hydrogen-bonding range, suggesting that one of the carboxylates is protonated in the ground state. To identify which residue is protonated and which serves as the primary counterion, we generated the E105Q and D231N mutants, which mimic protonated E105 and D231, respectively, and measured their absorption spectra at different pH values, comparing them with WT (Fig. 2d, Supplementary Fig. 3d, e). Notably, the *λ*_max_ of E105Q (574 nm) was nearly identical to that of WT (578 nm), whereas D231N showed a significant blue shift (546 nm). These results indicate that in WT ChR024 at neutral pH, E105 is protonated and D231 deprotonated, and that protonation of E105 makes a significant contribution to the red-shifted spectrum. Although C1C2 also carries Glu and Asp at these positions, its *λ*_max_ is much shorter (470 nm), likely because both residues are deprotonated in the ground state due to the presence of a nearby lysine (K132 in C1C2 and K93 in *Cr*ChR2)^38^. To test this idea, we substituted the corresponding residue in ChR024, Y80, with lysine. As expected, the Y80K mutant exhibited a *λ*_max_ (539 nm) similar to that of D231N, indicating that E105 becomes deprotonated in this mutant and that a non-basic residue at this position is critical for maintaining the high p*K*_a_ of E105. This conclusion is consistent with previous studies of Chrimson and *Ca*ChR1, other CCRs that also carries Glu and Asp on TM3 and TM7 and exhibits a red-shifted absorption spectrum^15,38,39^.

### Long-distance effect in color tuning via p*K*a modulation of the counterion

ChR024 was originally identified from a genomic database by ML-based screening focusing on sequences of 24 amino acids surrounding the retinal binding pocket^29^. This approach successfully extracted ChR024 as a ChR with a predicted *λ*_max_ of 543 nm, but a non-trivial gap remained compared with the experimentally determined value of 578 nm. We hypothesized that this gap might arise from additional residues important for color tuning that were not included in the original 24-residue set (Fig. 3a). To improve the ML-based *λ*_max_ prediction method and further our understanding of the color-tuning mechanism of ChR024, we next performed quantum mechanical/molecular mechanical (QM/MM) calculations (Methods).

**Fig. 3.**
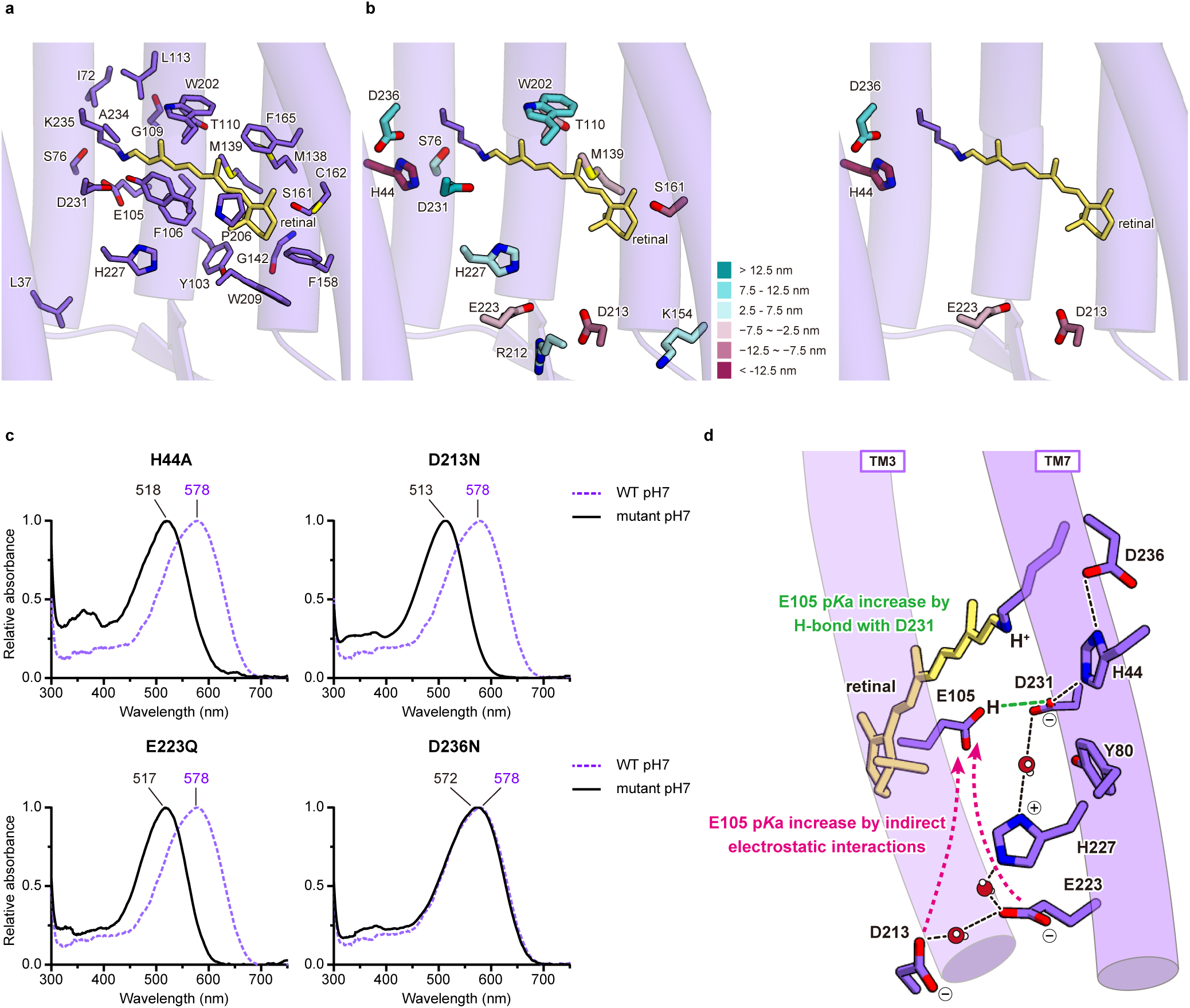
| Red color tuning mechanism of ChR024. **a,** Distribution of residues used for machine learning–based *λ*_max_ prediction. **b,** (left) Mapping of residues calculated by QM/MM to shift *λ*_max_ by more than 2.5 nm. Colors represent the magnitude of the effect, ranging from +12.5 nm (red) to –12.5 nm (teal). (right) Residues with predicted shifts greater than +5 nm or less than –5 nm that are located outside the region identified by machine learning. **c,** Relative absorption spectra of mutants predicted to have large effects: H44A (top left), D213N (top right), E223Q (bottom left), and D236N (bottom right) at pH 7.0. Each spectrum is normalized to the absorbance peak corresponding to retinal (set to 1.0). The absorption spectrum of WT ChR024 is shown as a dashed purple line. **d,** Schematic model of the red-shifted absorption mechanism in ChR024. Black lines indicate hydrogen bonds. The pink dashed arrow and green dashed line represent the long-range electrostatic effect and the hydrogen bond between counterions on TM3 and TM7, respectively. Water molecules are shown as red (oxygen) and white (hydrogen) spheres. TM3 and TM7 are illustrated as purple and light purple tubes, respectively, with other helices omitted for clarity. Retinal and key residues involved in red-shifted absorption are shown as sticks, colored yellow and purple, respectively.

Using the cryo-EM structure, we first calculated the protonation states of all residues in ChR024 and then, based on this information, evaluated the contributions of electrostatic interactions between individual residues and the retinal to the opsin shift, defined as the change in *λ*_max_ of the absorption spectrum. The calculations confirmed that E105 is protonated and D231 deprotonated (Supplementary Data 1), and identified 13 charged or polar residues that significantly affect the opsin shift (>2.5 nm) (Fig. 3b left, Supplementary Data 1). Notably, H44, K154, R212, D213, E223, and D236 were not included in the original 24-residue set, and four of these (H44, D213, E223, and D236) were predicted to contribute more than 5 nm to the *λ*_max_ shift (Fig. 3b right, Supplementary Data 1). We therefore generated mutants of these four residues (H44A, D213N, E223Q, and D236N) and measured their absorption spectra (Fig. 3c, Supplementary Fig. 3d, e).

As shown in Fig. 3c, all four mutants exhibited blue-shifted spectra (518, 513, 517, and 572 nm, respectively). However, the results were unexpected: the predicted opsin shifts for H44A, D213N, E223Q, and D236N based on the electrostatic contribution of each residue were –19, –10, –5, and +13 nm, respectively, whereas the observed shifts were –60, –65, –61, and –6 nm. Except for D236N, the mutational effects were far larger than predicted and cannot be explained solely by the loss of electrostatic interactions between the residues and the retinal. Moreover, the magnitudes of the opsin shifts were nearly identical for H44A, D213N, and E223Q. Because the *λ*_max_ values of these three mutants were similar to that of WT ChR024 at pH 9 (526 nm), a condition in which both E105 and D231 are deprotonated, we hypothesized that the ∼60 nm shift arises because these mutations lower the p*K*_a_ of E105, leading to its deprotonation (Supplementary Fig. 3d, e). To evaluate this hypothesis, we calculated the contribution of each residue’s electrostatic potential to the p*K*_a_ of E105 and found that D213 and E223 make relatively large contributions (+1.0 and +1.8, respectively). Considering that the p*K*_a_ of E105 estimated from pH titration experiments of WT ChR024 is 8.25^29^, it is reasonable to conclude that the D213N and E223Q mutations decrease the p*K*_a_ of E105 sufficiently to cause its deprotonation. In contrast, H44 is located close to E105, and the H-to-A mutation not only alters the electrostatic interaction but also reduces the side-chain volume. We therefore suggest that the H44A mutation creates extra space near E105, allowing water molecules to enter and trigger its deprotonation. Further details are provided in the Supplementary Notes. Collectively, these findings highlight that E105 protonation is governed not only by its short-range interaction with D231 but also by long-range electrostatics and local solvation, making it highly sensitive to even subtle perturbations that trigger deprotonation and a pronounced blue shift in the absorption spectrum (Fig. 3d).

### Ion-conducting pores

The overall position of ion-conducting pathways is known to be similar between channel- and pump-type rhodopsins, being located between TM1, TM2, TM3, TM6, and TM7 within each monomer^31,40^. To understand the mechanism of cation conduction in ChR024, we compared the structures of the ion-conducting pores in ChR024, *Hs*BR, ChRmine, C1C2, and archaerhodopsin-3 (AR3), another prototypical proton-pumping rhodopsin widely used as an optogenetic tool^16,18,41,42^ (Fig. 4).

**Fig. 4.**
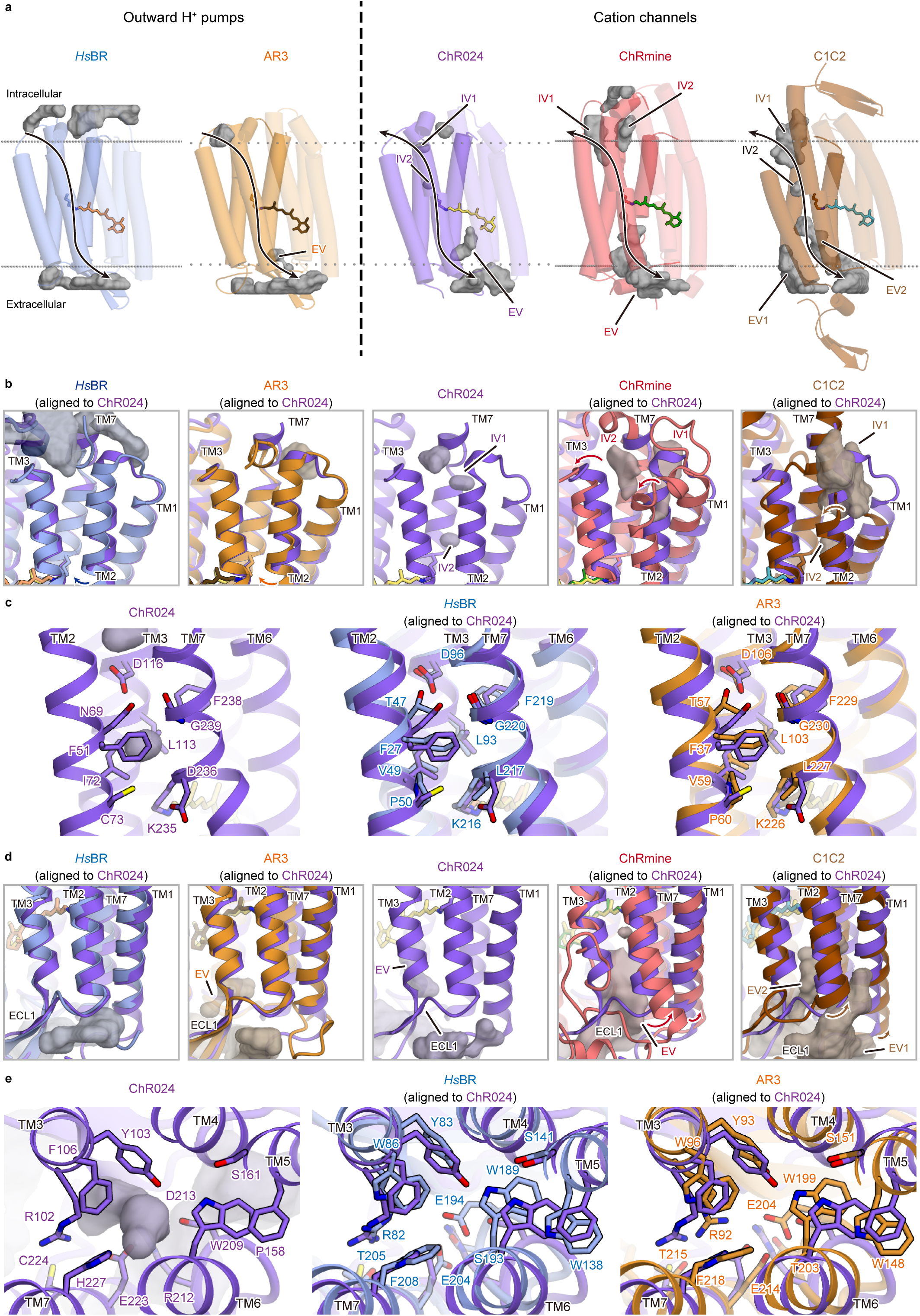
| Ion-conducting pathway. **a,** Comparison of ion-conducting pores in ChR024 (middle) with those of outward proton pump rhodopsins (*Hs*BR, blue; AR3, orange; left and second from left) and cation channelrhodopsins (ChRmine, red; C1C2, brown; second from right and right). Transmembrane helices are shown as tube models, with TM4, TM5, and TM6 in semitransparent colors. Intra- and extracellular cavity surfaces were calculated with HOLLOW^80^ and are shown semitransparently in dark gray. Gray dots represent membrane boundaries calculated with the TmDet 4.0 web server.^81^ Black arrows indicate putative ion conducting pathway. **b,** Architectures of ChR024 around intracellular cavity (middle) and comparison with *Hs*BR, AR3, ChRmine, and C1C2. Curved arrows indicate helical differences relative to ChR024. **c,** Magnified view of the intracellular side of ChR024 and its cavity (left), and comparisons with *Hs*BR (middle) and AR3 (right) viewed parallel to the membrane **d,** Architecture of ChR024 around the extracellular cavity (middle) and comparison with *Hs*BR, AR3, ChRmine, and C1C2. Curved arrows indicate helical differences relative to ChR024. **e,** Magnified view of the extracellular side of ChR024 and its cavity (left), and comparisons with *Hs*BR (middle) and AR3 (right) viewed from the intracellular side.

The structural comparison revealed that ChRmine and C1C2 contain large internal cavities, *Hs*BR and AR3 have almost none, and ChR024 falls in between (Fig. 4a). This is intriguing because previous studies have suggested that cavity size in the ground state is a key determinant of the molecular function of ion-translocating rhodopsins^43^: if the cavity is too small, the intra- and extracellular cavities never connect and the protein functions as an ion pump, whereas if the cavity is large enough to connect during the photocycle, the protein behaves as a leaky pump—in other words, an ion channel. ChR024 shares several structural features with ion-pumping rhodopsins (Figs. 1 and 2), but its cavity size lies between those of ion pumps and canonical ChRs.

Further analysis revealed that ChR024 has smaller intracellular cavities than ChRmine and C1C2, mainly due to the disposition of TM2, TM3, and/or ICL1 (Fig. 4b). In ChRmine, the intracellular sides of TM2 and TM3 are tilted outward, whereas in C1C2, TM2 is tilted and ICL1 is flexible; both arrangements create relatively large intracellular vestibules (IVs). By contrast, the intracellular regions of *Hs*BR and AR3 are tightly packed, leaving no large IVs. While the overall intracellular arrangement of TM1, TM2, TM3, and TM7 is similar among ChR024, *Hs*BR, and AR3, ChR024 shows a slight displacement of TM1 and TM7 and a proline residue (P50 in *Hs*BR and P60 in AR3) is replaced with a cysteine (C73) on TM2 (Fig. 4c). Together, these structural elements contribute to the formation of a narrow vestibule (IV2) near the Schiff base lysine (K235).

On the extracellular side, ChR024 again has smaller cavities than ChRmine and C1C2, mainly due to the disposition of TM1, TM2, and/or TM3 (Fig. 4d). In ChRmine, the extracellular sides of TM1 and TM2 are tilted outward and TM3 is partially unfolded, enlarging the cavity. In C1C2, the extracellular sides of TM1 and TM2 are also tilted outward, creating a relatively large extracellular vestibule (EV). By contrast, in ChR024, TM1, TM2, TM3, and TM7 superimpose well with those in *Hs*BR and AR3, but the last helical turn of TM6 and ECL3 adopts a unique conformation. This structure, together with the displacement of four aromatic residues (Y103/F106/W209/H227 in ChR024, Y83/W86/W189/F208 in *Hs*BR, and Y93/W96/W199/F218 in AR3), enlarges the cavity in ChR024 (Fig. 4e).

To gain further insight into the protein dynamics of ChR024, particularly how the channel opens and closes and how its gating mechanism compares with those of other channel- and pump-type rhodopsins, we performed Fourier-transform infrared (FTIR) spectroscopy on ChR024 and ChRmine, and compared the results with previously reported data for *Hs*BR^44–47^ (Fig. 5). Specifically, we identified the temperatures at which intermediates in the photocycle can be trapped (Fig. 5a, Supplementary Fig. 4a), measured the FTIR spectra at each temperature (Fig. 5b, Supplementary Fig. 4b), and evaluated conformational changes from the peak-pair region of the amide I signal (Fig. 5c, Methods). The results revealed that all three rhodopsins undergo large conformational changes during transitions from 230 K to 265/277 K, corresponding to the L3-to-N, M-to-N, and M1/M2-to-M2 transitions in ChR024, *Hs*BR, and ChRmine, respectively. It has been reported that Schiff base accessibility switches from the extracellular to the intracellular side during the M-to-N transition in *Hs*BR^31^, and laser patch-clamp experiments suggest that the channel pore opens during the L3-to-N and M1/M2-to-M2 transitions in ChR024 and ChRmine, respectively^29^ (manuscript in preparation). Taken together, these results suggest that although the timing (i.e., the specific intermediate) differs among rhodopsins, large structural rearrangements are consistently coupled to the key gating events in each protein: Schiff base accessibility switching in *Hs*BR, and channel opening in ChR024 and ChRmine (Fig. 5d).

**Fig. 5.**
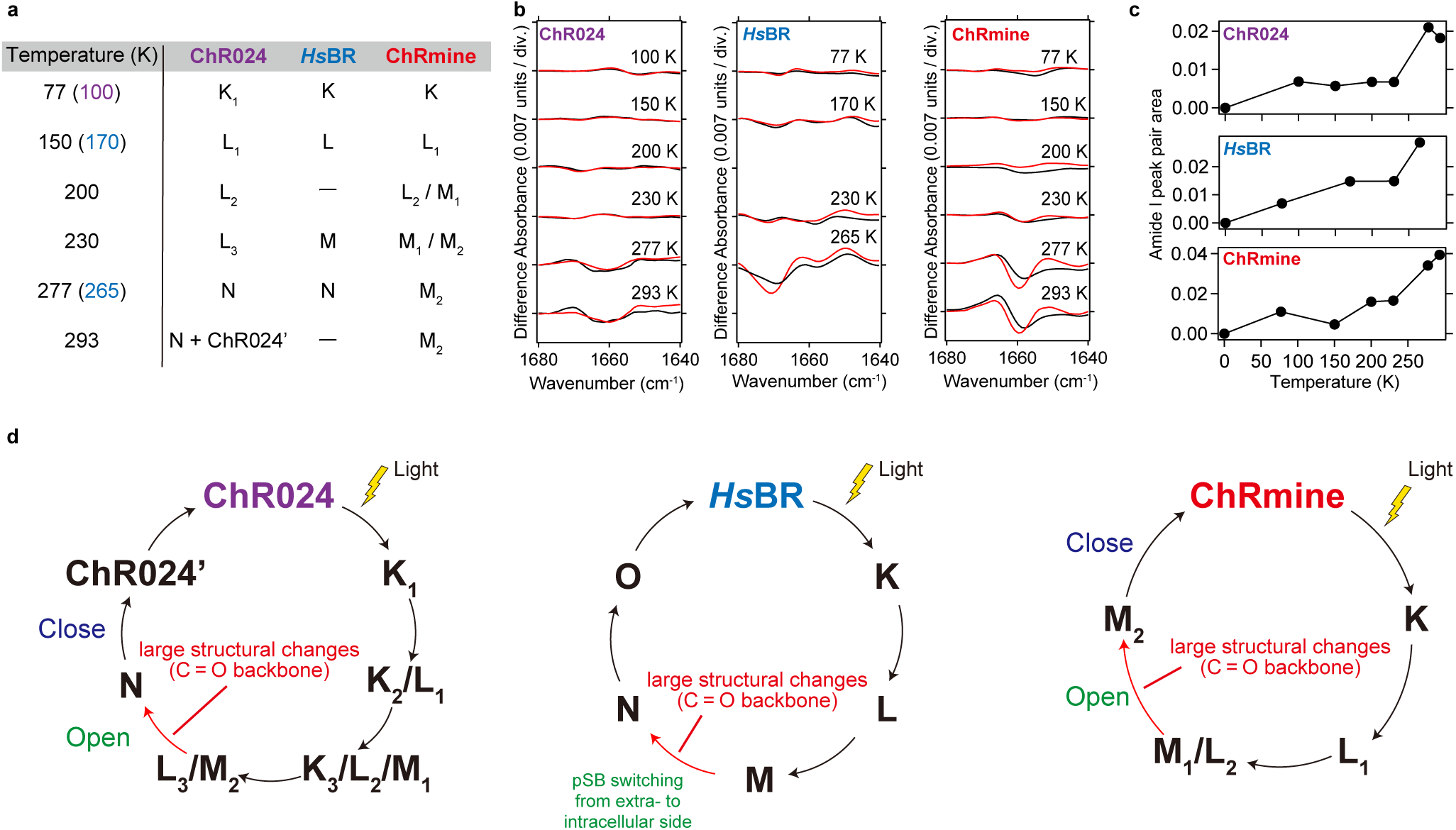
| Comparison of conformational dynamics of ChR024, *Hs*BR, and ChRmine. **a**, Comparison of intermediate accumulated at each temperature. **b,** Light-induced low-temperature difference FTIR spectra of ChR024 (left), *Hs*BR (middle), and ChRmine (right) in H_2_O (black) and D_2_O (red), in the 1680–1640 cm^-1^ region. The FTIR spectra of *Hs*BR were reproduced from Ref 44–47. **c,** Temperature dependence of the amide I peak area calculated from the light-induced low-temperature difference FTIR spectra in (**b**). **d,** Photocycle and conformational dynamics of ChR024, *Hs*BR, and ChRmine determined from changes in the amide I peak area based on (**c**).

### Functional conversion from pump to channel

Detailed structural comparisons revealed that among ion-pumping rhodopsins, ChR024 is more similar to AR3 than to *Hs*BR (Fig. 4). This observation motivated us to experimentally probe the functional boundary between channel and pump by attempting to convert AR3 from a light-driven proton pump into a light-gated ion channel. AR3 has a larger EV than *Hs*BR, though still smaller than that of ChR024 (Figs. 4a, 6a). To make the structure and properties of AR3 more similar to those of ChR024, we swapped three residues in the EV one by one between AR3 and ChR024 and evaluated their functions by measuring the reversal potential (*E*_rev_) (Fig. 6b–f).

**Fig. 6.**
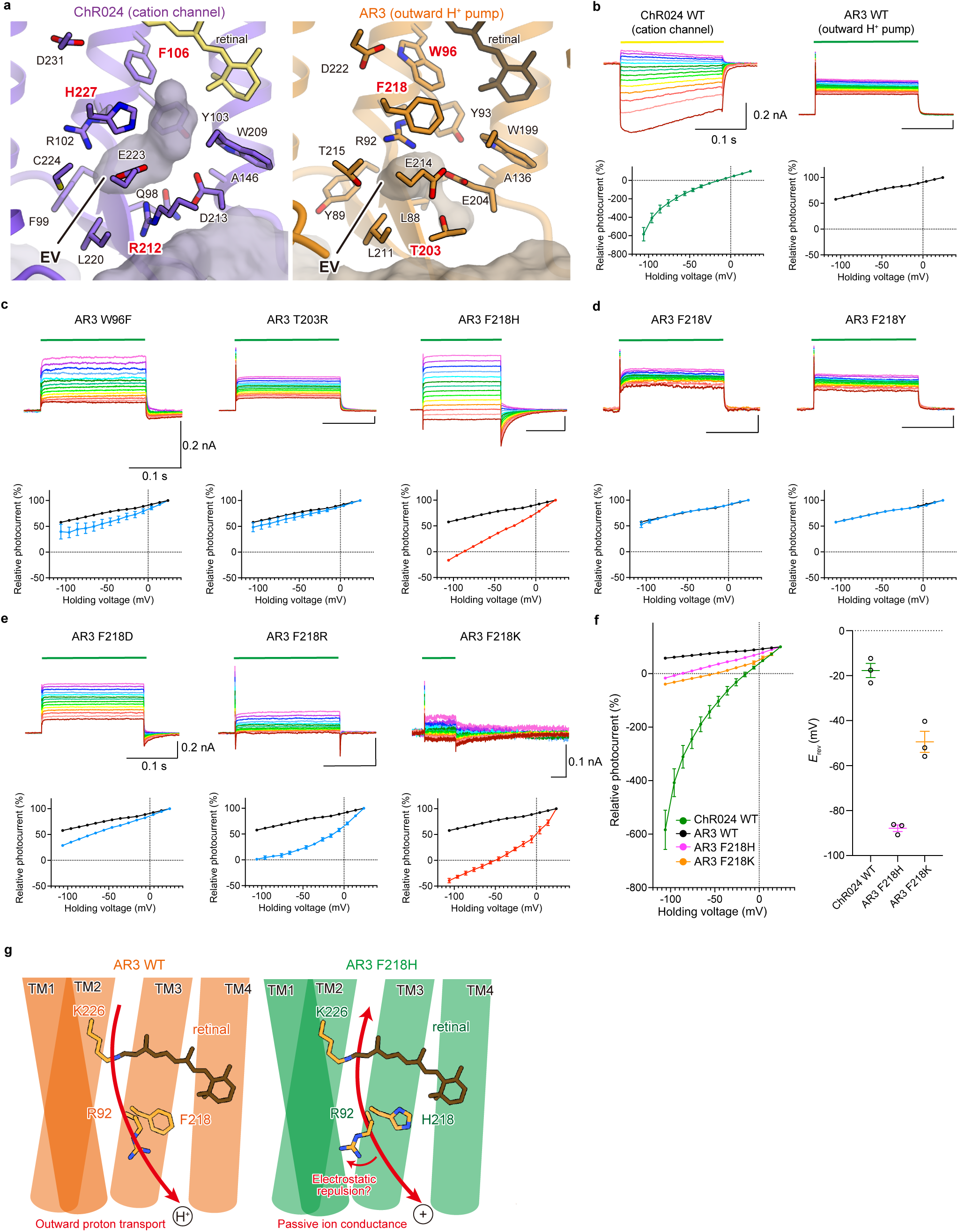
| Electrophysiological characterization of AR3 mutants. **a,** Comparison of amino acid residues around the EV between ChR024 and AR3. Residues swapped for functional conversion of AR3 are shown in red. **b,** Photocurrents (top) and *I*-*V* plots (bottom) of ChR024 WT and AR3 WT. Photocurrents in *I*-*V* plots are normalized photocurrents at 24 mV. Mean ± s.e.m. (*n* = 3). **c,** Photocurrents (top) and *I*-*V* plots (bottom) of AR3 mutants with residues swapped with ChR024 at W96, T203, and F218. Photocurrents in *I*-*V* plots are normalized at 24 mV. Mean ± s.e.m. (*n* = 3). For comparison, the *I*-*V* plot of AR3 WT (black curve) is shown in all panels. Mutant *I*-*V* plots are shown as red or blue curves. **d-e,** Photocurrents (top) and *I*-*V* plots (bottom) of AR3 F218 mutants. F218 was substituted with smaller or larger residues **(d)** or negatively or positively charged residues **(e)**. Photocurrents in *I*-*V* plots are normalized at 24 mV. Mean ± s.e.m. (*n* = 3). For comparison, the *I*-*V* plot of AR3 WT (black curve) is shown in all panels. Mutant *I*-*V* plots are shown as red or blue curves. **f,** Summary of *I*-*V* plots of ChR024 WT, AR3 WT, AR3 F218H, and AR3 F218K (left). Photocurrents in *I*-*V* plots are normalized at 24 mV. Mean ± s.e.m. (*n* = 3). Summary of reversal potentials (*E*_rev_) of ChR024 WT, AR3 F218H, and AR3 F218K (right). Mean ± s.e.m. (*n* = 3). **g,** Functional conversion model of the AR3 F218H mutant.

First, the *E*_rev_ of WT ChR024 was –17.7 ± 3.1 mV, whereas WT AR3 showed no measurable *E*_rev_ within the physiological range, consistent with the fact that WT ChR024 functions as a channel and WT AR3 as an ion pump (Fig. 6b). The same was true for the W96F and T203R mutants of AR3, which did not show an *E*_rev_ above –100 mV and retained potent proton-pumping activity (Fig. 6c). By contrast, the F218H mutant exhibited dramatically different behavior, with an *E*_rev_ of –87.9 ± 1.5 mV (Fig. 6c). Because an F-to-H substitution alters hydrophilicity, bulkiness, and charge simultaneously, we next tested five additional mutations at position F218 (Fig. 6d, e). The results showed that F218V and F218Y displayed almost identical current–voltage (*I*–*V*) curves to WT AR3 (Fig. 6d), whereas F218D, F218R, and F218K mutants significantly altered them (Fig. 6e). Among these, F218K showed the largest effect, with an *E*_rev_ as high as –49.4 ± 4.7 mV, suggesting that changes in the electrostatics of this residue play a key role in functional conversion (Fig. 6f).

Since F218 is located in close proximity to R92, it is likely that F218H and F218R introduce electrostatic repulsion with R92, thereby destabilizing the local interaction network and expanding the EV (Fig. 6g). Such changes may enable passive ion conduction through the ion-conducting pore.

## Discussion

Our structural, spectroscopic, and computational analyses of ChR024 suggest that the short distance between the counterion pair on TM3 and TM7, the absence of basic residues (lysine or arginine) on TM2, and even charged residues located far from the counterion (about 10–13 Å) are critically important for maintaining protonation of the counterion on TM3, thereby causing the significant red shift in the absorption spectrum (Fig. 3). This is consistent with previous studies of Chrimson and *Ca*ChR1, although they belong to distinct subfamilies of ChRs, suggesting that this principle may be broadly applicable to other ChRs. Intriguingly, a recent study of HulaChrimson, a newly discovered CCR with glutamate and aspartate as counterions (E184 and D314), reported that E184 is deprotonated despite the absence of basic residues on TM2 at the position corresponding to K132 in C1C2^48^. The structural basis for the deprotonated E184 and the resulting blue-shifted spectrum remains unclear. However, considering the long-distance effects on the counterion’s p*K*_a_ observed in ChR024, it is plausible that charged residues positioned distantly from E184 collectively lower its p*K*_a_ and trigger its deprotonation—the opposite case to ChR024.

These long-distance effects of charged residues should also be considered for accurate prediction of *λ*_max_. Indeed, electrostatic calculations of ChR024 reveal that all charged residues in the protein cavity influence the p*K*_a_ of the counterion by more than ∼1.0, provided they are not exposed to bulk solvent (Supplementary Data 1). (By contrast, residues exposed to solvent do not significantly affect the p*K*_a_ of the counterion because of their greater distance and the shielding effect of water.) Thus, with advances in protein structure prediction software^49^, integrating predicted structural information with ML-based methods will further enhance the gene mining of ChRs with desired spectral properties.

In this study, we also converted the function of AR3 from a proton pump to an ion channel by substituting the residue at position F218 with a basic amino acid. Similar functional conversions have been reported in KR2, *Cs*R, and *Ap*Ops1-3, and our findings with AR3 are consistent with these observations^50–53^. Mutations within the extracellular vestibule likely destabilize the interaction network centered on arginine (R102 in ChR024), loosen it, and lead to passive ion conductance at intermediates in which the extracellular gate should close to prevent leak current. To fully understand the mechanisms distinguishing channels from pumps, further studies will be needed, such as converting a channel-type rhodopsin into a pump. Nonetheless, the present study enhances our understanding of the structural basis for color tuning and ion transport in rhodopsins, sheds light on their evolutionary diversification, and provides a foundation for the rational design of next-generation optogenetics tools.

## Methods

### Expression and purification of ChR024

The wild-type ChR024 (residues M1–T280) was modified to include a C-terminal Kir2.1 membrane-targeting sequence, a human rhinovirus 3C protease cleavage site, enhanced green fluorescent protein (eGFP), and an 8×His tag, and cloned into the pFastBac vector. Constructs were expressed in *Spodoptera frugiperda* (Sf9) insect cells using the pFastBac baculovirus system. Sf9 cells were grown in suspension to a density of 3.0 × 10^6^ cells ml^−1^, infected with baculovirus, and shaken at 27.5 °C for 24 h. All-*trans* retinal (ATR) (Wako) was supplemented to a final concentration of 10 µM in the culture medium 24 h after the infection. Cells were harvested and resuspended in a hypotonic lysis buffer (20 mM HEPES-NaOH pH 7.5, 20 mM NaCl, 10 mM MgCl_2_, 1 mM benzamidine, 1 µg ml^−1^ leupeptin, 10 µM ATR). The cell suspension was collected by centrifugation at 8,000 ×g for 5 min, and this process was repeated twice. The cell pellets were then disrupted using with a glass dounce homogenizer in a hypertonic lysis buffer (20 mM HEPES-NaOH pH 7.5, 1 M NaCl, 10 mM MgCl_2_, 1 mM benzamidine, 1 µg ml^−1^ leupeptin, 10 µM ATR), and the crude membranes were collected by ultracentrifugation (45Ti rotor, 125,000 ×g for 1 h). The procedure was repeated twice, and the membranes were subsequently homogenized in membrane storage buffer (20 mM HEPES-NaOH pH 7.5, 500 mM NaCl, 10 mM imidazole, 20% glycerol, 1 mM benzamidine, 1 µg ml^−1^ leupeptin), flash-frozen in liquid nitrogen, and stored at −80 °C until use. For solubilization, the membrane fraction was incubated in a solubilization buffer (1% n-dodecyl-β-D-maltoside (DDM) (Glycon), 20 mM HEPES-NaOH pH 7.5, 500 mM NaCl, 20% glycerol, 10 mM imidazole, 1 mM benzamidine, 1 µg ml^−1^ leupeptin) at 4 °C for 2 h. The insoluble cell debris was removed by ultracentrifugation (45Ti rotor, 125,000 ×g for 1 h), and the supernatant was incubated with the Ni-NTA superflow resin (QIAGEN) at 4 °C for 2 h. The Ni-NTA resin was loaded onto an open chromatography column, washed with 2.5 column volumes of wash buffer (0.05% DDM, 20 mM HEPES-NaOH pH 7.5, 100 mM NaCl, and 50 mM imidazole) three times, and eluted with elution buffer (0.05% DDM, 20 mM HEPES-NaOH pH7.5, 100 mM NaCl, and 300 mM imidazole). After tag cleavage with His-tagged 3C protease, the sample was reapplied onto the Ni-NTA open column to remove the cleaved eGFP-His-tag and His-tagged 3C protease. The flow-through fraction was collected, and concentrated to approximately 2 mg ml^−1^ using an Amicon Ultra centrifugal filter unit (50 kDa cutoff) (Merck Millipore). The concentrated samples were ultracentrifuged (TLA55 rotor, 71,680 ×g for 30 min) before size-exclusion chromatography on a Superdex 200 Increase 10/300 GL column (Cytiva), equilibrated in SEC buffer (0.03% DDM, 20 mM HEPES-NaOH pH 7.5, 100 mM NaCl). The peak fractions of the protein were collected and concentrated to approximately 10 mg ml^−1^.

### Preparation of membrane scaffold protein

Membrane scaffold protein (MSP1E3D1) was expressed and purified as described earlier^27,54^. MSP1E3D1 gene in pET-43a (+) was transformed into *Escherichia coli* (*E. coli*) BL21 (DE3) cells. Cells were grown at 37 °C with shaking to an OD600 of 0.5–1.0, and expression of MSP1E3D1 was induced by addition of 1 mM IPTG. After 4 h of incubation at 37 °C, cells were harvested by centrifugation. The cell pellets were resuspended in PBS (–) buffer supplemented with 1% Triton X-100 and protease inhibitors, and lysed by sonication. The lysate was centrifuged at 30,000×g for 30 min, and the supernatant was applied to a Ni-NTA column preequilibrated with lysis buffer. The column was washed sequentially with four bed volumes each of wash buffer-1 (40 mM HEPES-NaOH pH 7.5, 300 mM NaCl, 1% Triton X-100), wash buffer-2 (40 mM HEPES-NaOH pH 7.5, 300 mM NaCl, 50 mM sodium cholate), wash buffer-3 (40 mM HEPES-NaOH pH 7.5, 300 mM NaCl), and wash buffer-4 (40 mM HEPES-NaOH pH 7.5, 300 mM NaCl, 20 mM imidazole), followed by elution with wash buffer-4 containing 300 mM imidazole.

The eluted MSP1E3D1 was dialyzed against buffer containing 10 mM HEPES-NaOH pH 7.5, 100 mM NaCl, and concentrated to approximately 10 mg ml^−1^ using an Amicon Ultra 10 kDa molecular weight cutoff centrifugal filter unit (Merck Millipore). The concentrated sample was clarified by ultracentrifugation (TLA55 rotor, 71,680 ×g for 30 min), and flashed-frozen in liquid nitrogen, and stored at –80 °C. Protein concentration was determined by measuring absorbance at 280 nm (extinction coefficient = 29,910 M^−1^ cm^−1^) measured by NanoDrop 2000c spectrophotometer (Thermo scientific).

### Nanodisc reconstitution of ChR024

Prior to nanodisc reconstitution, 30 mg SoyPC (Sigma P3644-25G) was dissolved in 500 µL chloroform, and dried under a nitrogen stream to form a lipid film. The residual chloroform was removed by overnight vacuum desiccation. The lipid film was rehydrated in buffer containing 1% DDM, 20 mM HEPES-NaOH pH 7.5, 100mM NaCl, and sonicated until the solution became clear, yielding a 10 mM lipid stock solution.

ChR024 was reconstituted into nanodiscs formed by the scaffold protein MSP1E3D1 and SoyPC at a molar ratio of 1:4:50 (monomer ratio: ChR024, MSP1E3D1, SoyPC). First, freshly purified ChR024 in SEC buffer (0.05% DDM, 20 mM HEPES-NaOH pH7.5, 100 mM NaCl) was mixed with SoyPC, and incubated on ice for 20 min. Purified MSP1E3D1 was then added to bring the total volume to 750 µL, and the mixture was gently rotated at 4 °C for 1 h. Detergents were removed by stepwise addition of Bio-Beads SM2 (Bio-Rad): 20 mg (26.7 mg mL^−1^) for the first 4 h, followed by 40 mg (53.3 mg mL^−1^) and 60 mg (80 mg mL^−1^) for two subsequent 4 h incubations at 4 °C. The sample was clarified by ultracentrifugation (TLA55 rotor, 71,680 ×g for 30 min) prior to size-exclusion chromatography on a Superdex 200 Increase 10/300 GL column (Cytiva), preequilibrated in buffer containing 20 mM HEPES-NaOH pH 7.5, and 100 mM NaCl. The peak fractions were collected and concentrated to approximately 16 mg ml^−1^ estimated from the absorbance at 280 nm (A_280_ = 30) using an Amicon Ultra 50 kDa molecular weight cutoff centrifugal filter unit (Merck Millipore).

### Cryo-EM grid preparation of DDM-solubilized and nanodisc-reconstituted ChR024

Prior to grid preparation, the freshly concentrated sample was clarified by ultracentrifugation (TLA55 rotor, 71,680 ×g for 30 min) at 4°C. Grids were glow-discharged with a PIB-10 plasma ion bombarder (Vacuum Device) at 10 mA current with the dial setting of 2 min on both sides. 3 µL aliquot of the protein solution was applied to freshly glow-discharged Quantifoil R1.2/1.3 Au 300 mesh holey carbon grid in a dark room under dim red light. Samples were vitrified by plunge-freezing into liquid ethane by liquid nitrogen using a FEI Vitrobot Mark IV (Thermo Fisher Scientific) at 4 °C with 100% humidity. The blotting force was set to 10, with a waiting time of 10 s and blotting time of 4 s.

### Cryo-EM data acquisition and image processing of DDM-solubilized ChR024

Cryo-EM images were acquired at 300 kV on a Titan Krios G3i microscope (Thermo Fisher Scientific) equipped with a Gatan BioQuantum energy filter and a K3 direct detection camera operated in electron-counting mode. The movie dataset was collected in standard mode, using the fringe-free imaging (FFI) and aberration-free image shift (AFIS) strategy in the EPU software (Thermo Fisher Scientific), with a nominal defocus range of −0.8 to −1.6 µm. The 2,136 movies were acquired at a dose rate of 13.8 e^-^/pixel/s, at a pixel size of 0.83 Å and a total dose of 46 e^-^/Å^2^.

Data processing was performed using the cryoSPARC v4.2.0 to v4.7.1 software packages^55^. The 2,136 movies were subjected to patch motion correction and patch CTF estimation in cryoSPARC. Initial particles were picked from all micrographs using blob picker and were extracted using a box size of 280 pixels. After 2D classification, 151,489 particles were selected from 557,139 particles. Two rounds of *ab initio* reconstruction and non-uniform refinement^56^ yielded the 3.62 Å map (C3 symmetry) from 65,293 particles. Further particles were picked using template picker, subjected to 2D classification and ab-initio reconstruction. Duplicates between the blob- and template-picked sets were removed before non-uniform refinement. Non-uniform refinement of the resulting dataset produced a final map at 3.22 Å global resolution (C3 symmetry) from 342,264 particles.

### Cryo-EM data acquisition and image processing of nanodisc-reconstituted ChR024

Cryo-EM images were acquired at 300 kV on a Titan Krios G3i microscope (Thermo Fisher Scientific) equipped with a Gatan BioQuantum energy filter and a K3 direct detection camera operated in electron-counting mode. Movies were collected in Correlated Double Sampling (CDS) mode using the fringe-free imaging (FFI) and aberration-free image shift (AFIS) strategy in the EPU software (Thermo Fisher Scientific), with a nominal defocus range of −0.8 to −1.6 µm. A total of 5,743 movies were acquired at a dose rate of 13.5 e^-^/pixel/s, at a pixel size of 0.83 Å and a total dose of 45.2 e^-^/Å^2^.

Data processing was performed using the cryoSPARC v4.2.1 to v.4.7.1 software packages. A total of 5,199,514 particles were picked using the template picker, subjected to two rounds of 2D classification and five rounds of ab initio refinement, yielding a 2.75 Å map (C3 symmetry) with 375,627 particles. Those particles were then used to train the Topaz picker^57^, resulting in the selection of 285,083 particles. Duplicates between the Topaz-picked particles (after 2D classification) and the best particles from template picking were removed, leaving 560,918 particles. These particles were subsequently subjected to ab initio reconstruction, non-uniform refinement, reference-based motion correction, non-uniform refinement, and local refinement with a TM mask, yielding a final map at 2.45 Å global resolution (C3 symmetry) with 415,817 particles.

### Model building and refinement

An initial model of ChR024 in nanodisc was generated by rigid body fitting of the predicted models of ChR024, generated using locally installed AlphaFold2^58^. This starting model was then subjected to iterative rounds of manual and automated refinement using Coot^59^ and Refmac5^60^ in Servalcat pipeline^61^, respectively. The Refmac5 refinement was performed with the constraint of C3 symmetry.

The final model was visually inspected for general fit to the map, and geometry was further evaluated using Molprobity^62^. The final refinement statistics are summarized in Table S1. All molecular graphics figures were prepared with UCSF Chimera^63^, UCSF ChimeraX^64^, and CueMol2 (http://www.cuemol.org).

### UV-Vis spectroscopy

To investigate the pH dependence of the UV-vis absorption spectra of WT ChR024, and its mutants (E105Q, D231N, Y80K, H44A, D213N, E223Q, and D236N), purified protein solution (Abs_280_ = 2-10) was 80-fold diluted in buffer containing 0.03% DDM, 100 mM NaCl, 100 mM of either citric acid pH 2.2, citric acid pH 3.0, sodium acetate pH 4.0, sodium citrate pH 5.0, sodium cacodylate pH 6.0, HEPES-NaOH pH 7.0, Tris-HCl pH 8.0, N-cyclohexyl-2-amino-ethanesulfonic acid (CHES) pH 9.0, 3-(cyclohexylamino)-1-propanesulfonic acid (CAPS) pH 10.0, or CAPS pH 11.0. The StockOptions pH Buffer Kit (Hampton research) was used for buffer preparation except for CHES pH 9.0 (Nacalai). The absorption spectra were measured with a V-750 UV-visible spectrometer (JASCO) at room temperature.

### Structure preparation for QM/MM analysis

The cryo-EM structure was used as the initial structure. The structure with R102 in the “A” conformation was analyzed. All hydrogen atom positions were energetically optimized using the CHARMM program^65^, with all heavy atoms fixed and all acidic and basic groups ionized in this procedure. Atomic charges and force field parameters were obtained from the CHARMM22 parameter set^66^. Atomic charges of the retinal Schiff base were determined by fitting the electrostatic potential of an isolated retinal Schiff base using the restrained electrostatic potential (RESP) procedure^67^. The C_β_ atom of the lysine was replaced with a methyl group. The electronic wave functions were calculated after geometry optimization. The restricted density functional theory (DFT) method was employed with the B3LYP functional and the LACVP* basis set, using the Jaguar program^68^.

### Protonation pattern and p*K*_a_

The protonation pattern of the protein was determined using the electrostatic continuum model, solving the linear Poisson-Boltzmann equation with the MEAD program^69^. All computations were performed at 300 K and pH 7.0, with an ionic strength of 100 mM. The experimentally measured p*K*_a_ values used as references were 12.0 for Arg, 4.0 for Asp, 9.5 for Cys, 4.4 for Glu, 10.4 for Lys, 9.6 for Tyr^70^, and 6.6 and 7.0 for the N_δ_ and N_ε_ atoms of His, respectively^71–73^. The dielectric constants were set to 4 for the protein interior and 80 for water. All water molecules were considered implicitly. The linear Poisson-Boltzmann equation was solved using a three-step grid-focusing procedure at resolutions of 2.5, 1.0, and 0.3 Å. The protonation pattern was sampled using the Monte Carlo method with the Karlsberg program^74^. The p*K*_a_ value of the focusing residue was evaluated by applying a bias potential to the residue to equalize the protonated and deprotonated populations ([protonated] = [deprotonated]); the resulting bias potential corresponds to the p*K*_a_ value. The electrostatic contribution of each side-chain to the p*K*_a_ value of the focusing residue was obtained from the shift in the p*K*_a_ value upon removal of the atomic charges of each side-chain.

### QM/MM calculation

The geometry was optimized using a QM/MM approach. The restricted DFT method was employed with the B3LYP functional and the LACVP* basis set, using the QSite program^75,76^. The QM region was defined as the retinal and the side-chain of Lys235 (Schiff base). All atomic positions in the QM region were fully optimized. In the MM region, all hydrogen atom positions were optimized using the OPLS2005 force field^77^ while the heavy atoms were fixed. The protonation pattern of titratable residues in the MM region was implemented in the atomic partial charges.

Based on the QM/MM-optimized geometry, the lowest excitation energy was calculated employing the polarized continuum model (QM/MM/PCM). In this model, polarization points were positioned on spheres with a radius of 2.8 Å from the center of each atom to account for possible water molecules in the cavity. A dielectric constant of 78.39 was applied. The restricted time-dependent DFT (TD-DFT) method was employed with the B3LYP functional and the 6-31G* basis set, using the GAMESS program^78^. The QM region and atomic partial charges in the MM region were set the same as in the geometry optimization. The calculated excitation energy (*E*_TD-DFT_) was corrected to obtain the absorption energy (*E*_abs_) using the following empirical equation^79^.

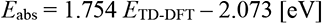

The resulting *E*_abs_ value was converted into the maximum absorption wavelength (*λ*_max_). The electrostatic contribution of each side-chain in the MM region to the absorption wavelength of the retinal was obtained from the shift in *λ*_max_ upon removal of the atomic charges of each side-chain.

### Pore analysis

Ion-conducting pore pathways were calculated with HOLLOW using a grid spacing of 1.0 Å^80^. Membrane boundaries were determined with the TmDeT 4.0 web server using default settings^81^. For both calculations, trimer structures were used as input. For AR3, monomer models were aligned to each protomer of the BR trimer (PDB ID: 5ZIM)^42^.

### Lipid reconstitution of ChR024 and ChRmine

As a membrane protein, rhodopsin requires reconstitution into a lipid environment to preserve its native behavior. Protein concentration was determined using a UV-visible spectrometer, and the optical density (O.D.) at the absorption maximum was adjusted to ∼0.5-1.0 by dilution with buffer (20 mM HEPES-NaOH, pH 7.5, 100 mM NaCl, 0.03% DDM). Soybean phospholipid (asolectin; Sigma-Aldrich) was added at a protein-to-lipid molar ratio of 1:20, and the mixture was incubated at 4 °C with gentle rotational mixing. Detergent removal and subsequent reconstitution into lipids were achieved by addition of Bio-Beads SM-2 (Bio-Rad). The reconstituted samples were repeatedly washed with buffer (2 mM K2HPO4/KH2PO4, pH 7.5) and pelleted by ultracentrifugation at 222,000 × g for 20 min at 4 °C in the dark.

### Low temperature UV-Visible spectroscopy

The reconstituted pellet was resuspended in buffer (2 mM K_2_HPO_4_/KH_2_PO_4_, pH 7.5) to a final concentration of 2.5 mg mL^-1^. A 60 μL aliquot was deposited onto a BaF_2_ window and dried with an aspirator. Prior to measurements, the films were hydrated with H_2_O, mounted on a sample holder, and placed in a cryostat (Optistat DN2, Oxford Instruments) connected to a UV-visible spectrometer (V750, JASCO).

Samples were cooled between 77 K and 230 K using liquid nitrogen, with the temperature controlled to ± 0.1 K. To generate photointermediates, ChR024 was illuminated for 2 min at 100 K, 150 K, 200 K, and 230 K with 570 nm, >500 nm, >500 nm, and >500 nm light, respectively, delivered from a 1-kW tungsten-halogen projector lamp (Rikagaku, Japan) through appropriate interference and cut-off filters (KL57, Y52; Toshiba). The K intermediate was reverted to the dark state at 100 K by >610 nm light (R63 cut-off filter, Toshiba), whereas the L intermediates were reverted at 150-230 K by 480 nm light (Toshiba interference filter).

Similarly, ChRmine was illuminated for 2 min at 77 K, 150 K, 200 K, and 230 K with 520 nm, >500 nm, >500 nm, and >500 nm light, respectively, using the same lamp and filters (KL52, Y52; Toshiba). The K intermediate was reverted to the dark state at 100 K by >550 nm light (O57 cut-off filter, Toshiba). The L intermediate at 150 K was reverted by 460 nm light (KL46 interference filter), and the M intermediates at 200 K and 230 K were reverted by 400 nm light (KL40 interference filter).

### Low temperature FTIR difference spectroscopy

Low-temperature Fourier-transform infrared (FTIR) spectroscopy was performed as previously described^82,83^. Light-induced changes of ChR024 and ChRmine were recorded using a transmission-type FTIR spectrometer (Cary 670, Agilent) equipped with a cryostat (Oxford Instruments). Film samples for FTIR measurements were prepared as described above for low-temperature UV-Vis spectroscopy. The optical density of the amide I band was adjusted to ∼0.8, and the films were hydrated with 1.5 μL of H_2_O, D_2_O, or D_2_^18^O before measurements.

For ChR024, intermediates were generated by illumination for 2 min at 100 K, 150 K, 200 K, and 230 K with 570 nm, >500 nm, >500 nm, and >500 nm light, respectively, using a 300 W xenon lamp (Max-303, Asahi Spectra) with appropriate interference and cut-off filters (KL57, Y52; Toshiba). The K intermediate at 100 K was reverted to the dark state by >610 nm light (R63 cut-off filter, Toshiba), whereas the L intermediates at 150-230 K were reverted by 480 nm light (Toshiba interference filter). For ChRmine, intermediates were generated at 77 K, 150 K, 200 K, and 230 K by illumination with 510 ± 10 nm (KL51 interference filter), >550 nm (O57 cut-off filter), >500 nm (Y52 cut-off filter), and >500 nm (Y52 cut-off filter) light, respectively, using a xenon lamp (LAX-103, Asahi Spectra). Illumination times were 1, 3, 5, and 5 min, respectively. The intermediate at 77 K was reverted to the dark state by >570 nm light (O59 cut-off filter, Toshiba) for 1 min.

For each FTIR measurement, interferograms were accumulated at a spectral resolution of 2 cm^-1^. For ChR024, the number of accumulated scans was as follows: 100 (H_2_O) and 80 (D_2_O) at 100 K; 40 (H_2_O) and 40 (D_2_O) at 150 K; 16 (H_2_O) and 8 (D_2_O) at 200 K; and 40 (H_2_O) and 40 (D_2_O) at 230 K. For ChRmine, 120, 240, and 240 interferograms were acquired in H_2_O, D_2_O, and D_2_^18^O at 77 K, respectively; 16 interferograms in each solvent at 150 K; 15 interferograms in each solvent at 200 K; and 15, 20, and 15 interferograms in H_2_O, D_2_O, and D_2_^18^O, respectively, at 230 K. The FTIR spectra of *Hs*BR were reproduced from Refs^44–47^.

The total peak area of the amide I band (1640-1680 cm^-1^) was calculated by first extracting peaks within this region and determining their average areas. The area of each peak was then estimated by multiplying the average peak area by its corresponding wavenumber range. The sum of these values was taken as the total amide I peak area.

### ATR-FTIR difference spectroscopy

Light-induced structural changes in the late intermediates of ChR024 and ChRmine were analyzed by attenuated total reflection (ATR)-FTIR spectroscopy, following a previously described protocol with minor modifications^27^. Reconstituted samples were placed on a silicon ATR crystal (Smiths, three internal reflections) and dried naturally, then rehydrated with 5 mL of buffer (200 mM NaCl, 200 mM Tris-HCl, pH 7.5).

Because the ATR-FTIR system was originally optimized for ion perfusion-induced difference spectroscopy using solution exchange, modifications were implemented to enable light-irradiation experiments. Specifically, a light source was installed above the ATR prism, with an optical interference filter (KL52, Toshiba) and a condenser lens positioned beneath the source. To record light-induced difference spectra, samples were illuminated for 5 min, followed by a 10 min dark adaptation, and this cycle was repeated three times. Spectral correction involved subtraction of contributions from unbound salts and water vapor, as well as baseline drift caused by protein swelling or shrinkage, yielding the final light-induced difference spectra.

### Cell culture and transfection for electrophysiology

ND7/23 cells were grown in Dulbecco’s modified Eagle’s medium (DMEM, Nacalai tesque) supplemented with 10% fetal bovine serum (FBS, Sigma-Aldrich) under a 5% CO_2_ atmosphere at 37°C. ND7/23 cells were seeded onto a collagen-coated 12-mm coverslips (4912-010, IWAKI) placing in a 4-well cell culture plate (30004, SPL Life Sciences). The expression plasmids (pCDNA3.1(+) vector encoding C-terminal mEYFP fused ChR024 or AR3 variants) were transfected in ND7/23 cells using polyethylenimine Max solution (Polysciences). 6-8 hours after the transfection, the medium was replaced by DMEM containing 10% horse serum, 50 ng/mL nerve growth factor-7S (Sigma-Aldrich), 1 mM N6,2’-O-dibutyryladenosine-3’,5’-cyclic monophosphate sodium salt (Nacalai tesque), and 1 μM Cytosine-1-β-D(+)-arabinofuranoside (FUJIFILM Wako Pure Chemical Co.). Electrophysiological recordings were conducted 2 days after the transfection.

### Electrophysiology

ND7/23 cells transfected with pcDNA3.1(+) plasmids were placed in a 295-305 mOsm extracellular solution (140 mM NaCl, 4 mM KCl, 2 mM CaCl_2_, 2 mM MgCl_2_, 10 mM HEPES pH 7.4, and 5 mM glucose). Borosilicate patch pipettes (Harvard Apparatus) with resistance of 2-4 MOhm were filled with 280-290 mOsm intracellular solution (120 mM potassium-gluconate, 10 mM EGTA, 2 mM MgCl_2_ and 10 mM HEPES pH 7.2). Light was delivered with the Spectra X Light engine (Lumencor) connected to the fluorescence port of a BX51WI microscope (Evident) with 575 ± 25 nm, and 550 ± 15 nm filter for yellow and green light generation. Photocurrents were measured in voltage clamp mode from -106 mV to 24 mV holding potential with 10 mV spacing (after liquid junction potential correction). For each recording, ChR024 transfected cells were illuminated with yellow (575 ± 25 nm) light, whereas AR3 and its variants were illuminated with green (550 ± 15 nm) light, each for 200 ms. The light power of yellow (575 ± 25 nm) light, and green (550 ± 15 nm) light was 203, and 163 mW/mm^2^. The obtained traces were baseline-corrected using the trace before light illumination by Igor Pro, and the traces from three recordings were averaged for *I*–*V* plot analysis. In the *I*–*V* plots, the photocurrent values were determined as the mean stationary current during the 200 ms light illumination, and the values were normalized to the photocurrent at +24 mV and plotted as a function of voltage. The reversal potential was determined by linear fitting of three data points around the zero-crossing of the photocurrent, and the x-intercept of the fitted line was taken as the reversal potential. For the functional analysis, statistical analyses were performed using the GraphPad Prism 10 (Ver 10.1.0) software (GraphPad), and the methods are described in the legends of the figures.

## Data Availability

The raw images of ChR024 in detergent micelles and lipid nanodiscs before motion correction have been deposited in the Electron Microscopy Public Image Archive under accession EMPIAR-xxxxx and EMPIAR-xxxxx. The cryo-EM density map and atomic coordinates for ChR024 in detergent micelle and lipid nanodiscs have been deposited in the Electron Microscopy DataBank: EMD-xxxxx and EMD-xxxxx, and PDB under accessions: xxxx and xxxx, respectively. All other data are provided in this article, its Supplementary Information and Source Data file, or from the corresponding author on reasonable request.

## Acknowledgements

We thank K. Hasegawa and A. Ohira (UTokyo) for administrative support. We also acknowledge ChatGPT, a multimodal large language model created by OpenAI, for providing guidance to improve the readability of this manuscript. Note that, after using this tool, we reviewed and edited the content as needed and took full responsibility for the content of the publication.

This work was supported by MEXT Promotion of Development of a Joint Usage/ Research System Project: Coalition of Universities for Research Excellence Program (CURE) (JPMXP1323015482 to K.I.), JSPS KAKENHI (25KJ0896 to Y.T., 24KJ0788 to S.T., JP22KJ1109 to M.T., JP24H02262 to M.F., JP23H02444 to H.Ishikita, JP24H02268/JP25H00424 to K.I., and 19H03163/22H00400/25H01338 to H.E.K.), AMED (24bm1123057h0001 to H.E.K.), JST PRESTO (JPMJPR24OF to M.F.), JST FOREST (JPMJFR204S to H.E.K.), and JST CREST (JPJPMJCR22N2 to K.I. and JPMJCR21P3/JPMJCR23B1 to H.E.K.).

## Author Contributions

Y.T., S.T., K.E.K., M.F., S.N., and M.W. performed cryo-EM analysis of ChR024. Y.T., S.T., and S.K. measured UV-Vis absorption spectra of ChR024 and its mutants. Y.T. and S.K. performed electrophysiological analysis. M.T. performed QM/MM calculations, supervised by H.Ishikita. Y.I., M.S., and Y.Y. performed FT-IR analysis, supervised by K.K., Y.F., and H.K. H.Ikeda conducted cloning and mutagenesis. K.I. provided input on structural considerations. Y.T., S.T., S.K., M.T., and H.E.K. wrote the manuscript with input from all authors. H.E.K. supervised all aspects of the research.

---

**Supplementary Fig. 1.**
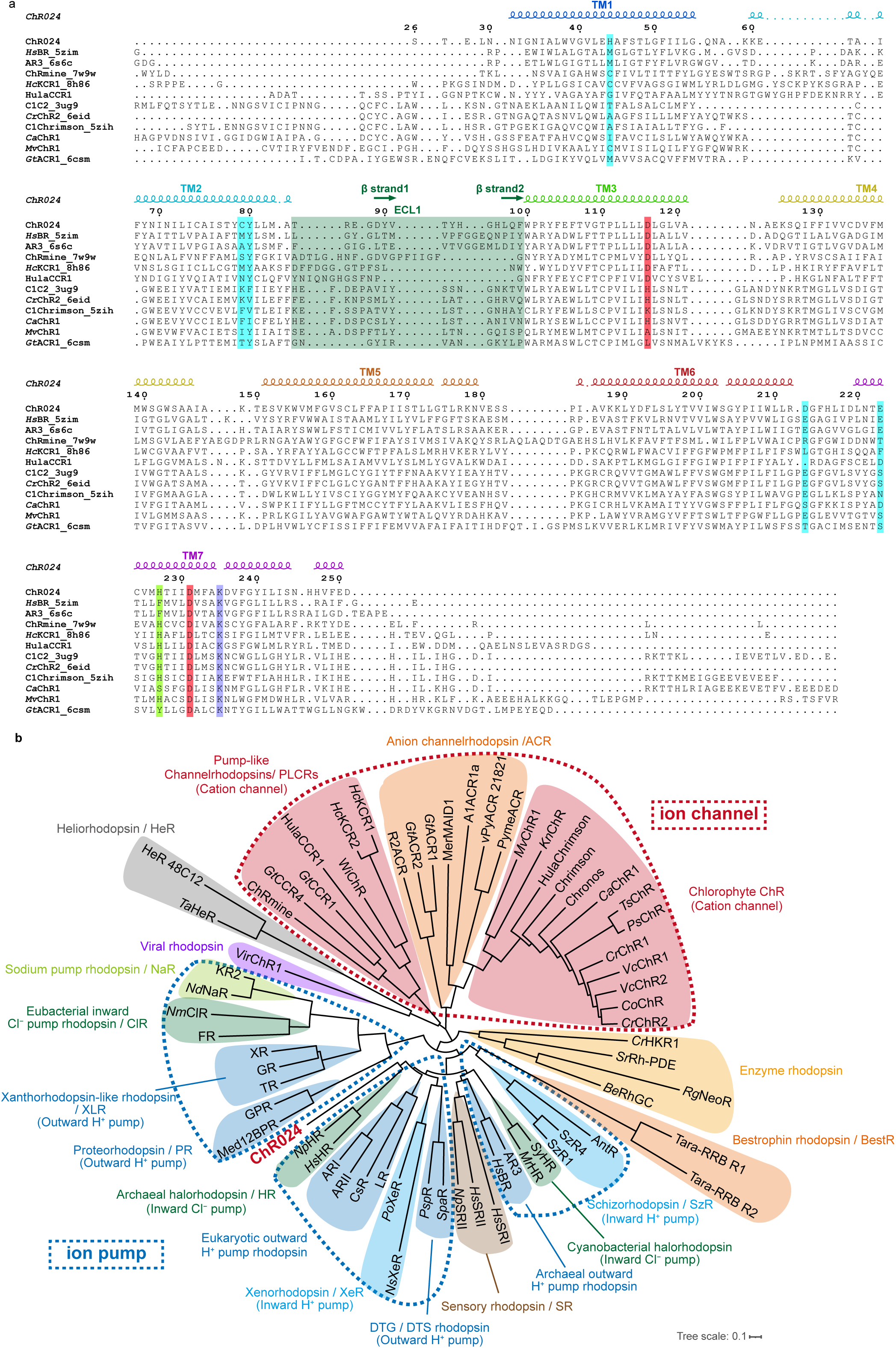
| Structure-based sequence alignment and phylogenetic analysis of microbial rhodopsins. a, Structure-based sequence alignment of ChR024 (GenBank: IMGM3300027749|Ga0209084_ 1026585|Ga0209084_10265853), *Hs*BR (PDB: 5ZIM)^42^, AR3 (PDB: 6S6C)^41^, ChRmine (PDB: 7W9W)^16^, *Hc*KCR1 (PDB: 8H86)^27^, HulaCCR1,^84^ the chimeric channelrhodopsin C1C2 (PDB: 3UG9)^18^, C1Chrimson (PDB: 5ZIH)^15^, *Cr*ChR2 (PDB: 6EID)^36^, *Ca*ChR1 (GenBank: AER58220.1), *Mv*ChR1 (GenBank: AEI83869.1), and *Gt*ACR1 (PDB: 6CSM)^85^. The alignment was generated using US-align^86,87^ and ESPript3 servers^88^. The input models for *Ca*ChR1, *Mv*ChR, and HulaCCR1 were generated using the AlphaFold3 web server.^89^ α-helical regions of ChR024 are shown as coils, and β-strands as arrows. The Schiff-base–bound lysine is highlighted in purple, counterion residues in red, key residues for red-shifted absorption in blue, ECL1 in green, and the critical residue for pump-to-channel conversion in light green. The alignment is shown starting from the N-terminus at the point where the gap rate is less than 80%. **b,** Phylogenetic tree of microbial rhodopsins. Each clade is colored according to function or clade designation, with ChR024 highlighted in red. Ion channels and ion pumps are indicated by red and blue dashed outlines, respectively.

**Supplementary Fig. 2.**
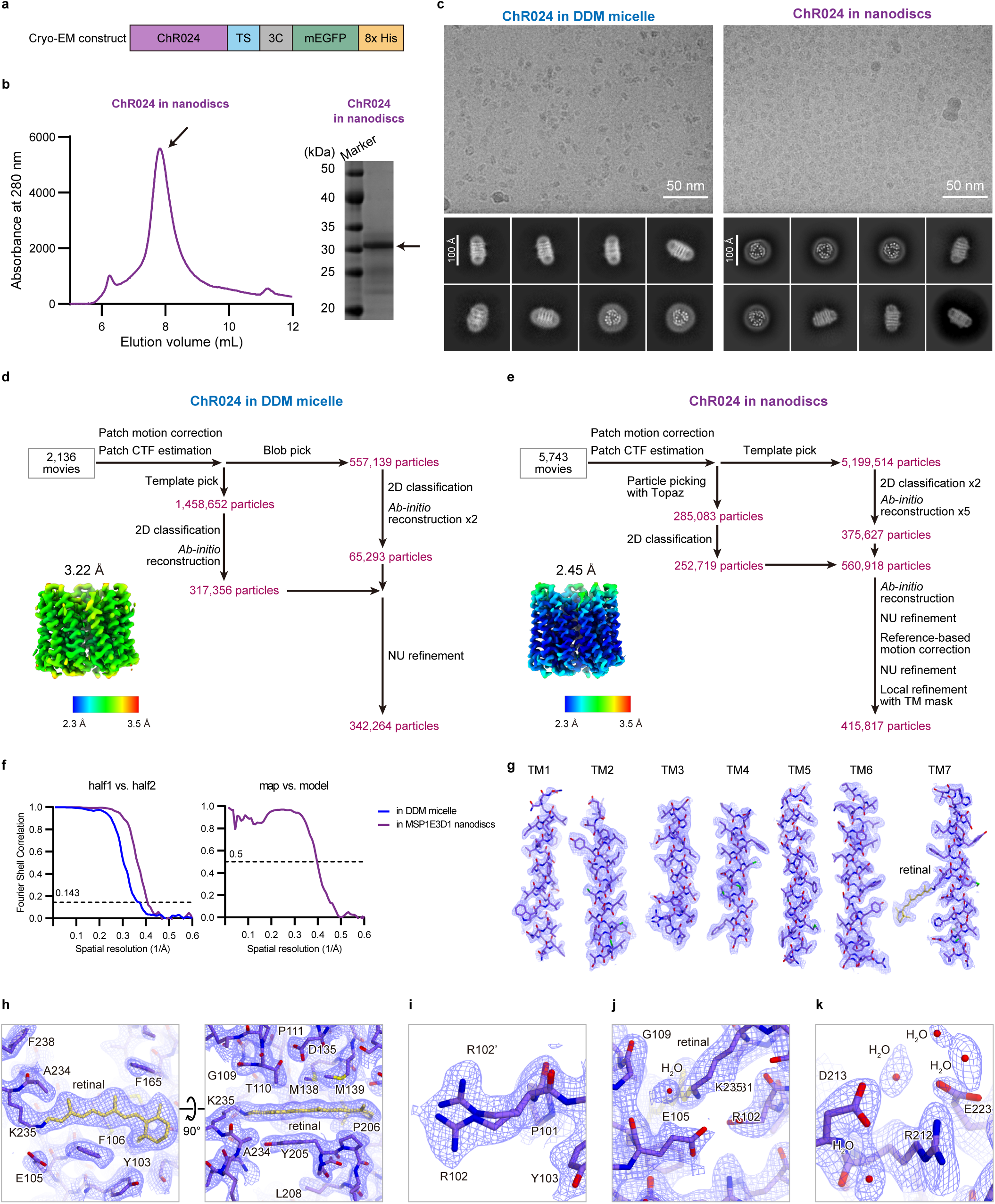
| 3D reconstruction of ChR024 in DDM micelles and MSP1E3D1 nanodiscs. **a**, Schematic representation of the ChR024 construct used for cryo-EM analysis. **b,** Representative SEC trace of ChR024 reconstituted in MSP1E3D1 nanodiscs with SDS-PAGE shown as an inset. The absorption signal was monitored by a UV/Vis detector at 280 nm. **c,** Representative cryo-EM micrographs of ChR024 in DDM micelles (top left) and in MSP1E3D1 nanodiscs (top right). Representative 2D class averages of ChR024 in DDM micelles (bottom left) and in MSP1E3D1 nanodiscs (bottom right). **d, e,** Data processing workflows for ChR024 in DDM micelles **(d)** and in MSP1E3D1 nanodiscs **(e)**. Final cryo-EM maps are colored according to local resolution (bottom left) **f,** Fourier shell correlation (FSC) between the two independently refined half-maps (left) and between the model and the map calculated from the model refined against the full reconstruction (right) of ChR024 in DDM micelles (blue) and in MSP1E3D1 nanodiscs (purple) **g–k,** Cryo-EM densities (blue mesh) and atomic models of ChR024: transmembrane helices **(g)**; retinal-binding pocket viewed parallel to the membrane (left) and from the intracellular side (right) **(h)**; R102 showing alternative conformations **(i)**; Schiff base region **(j)**; and extracellular region around D213 and E223 **(k)**. Stick models are shown for selected residues (purple) and retinal (yellow).

**Supplementary Fig. 3.**
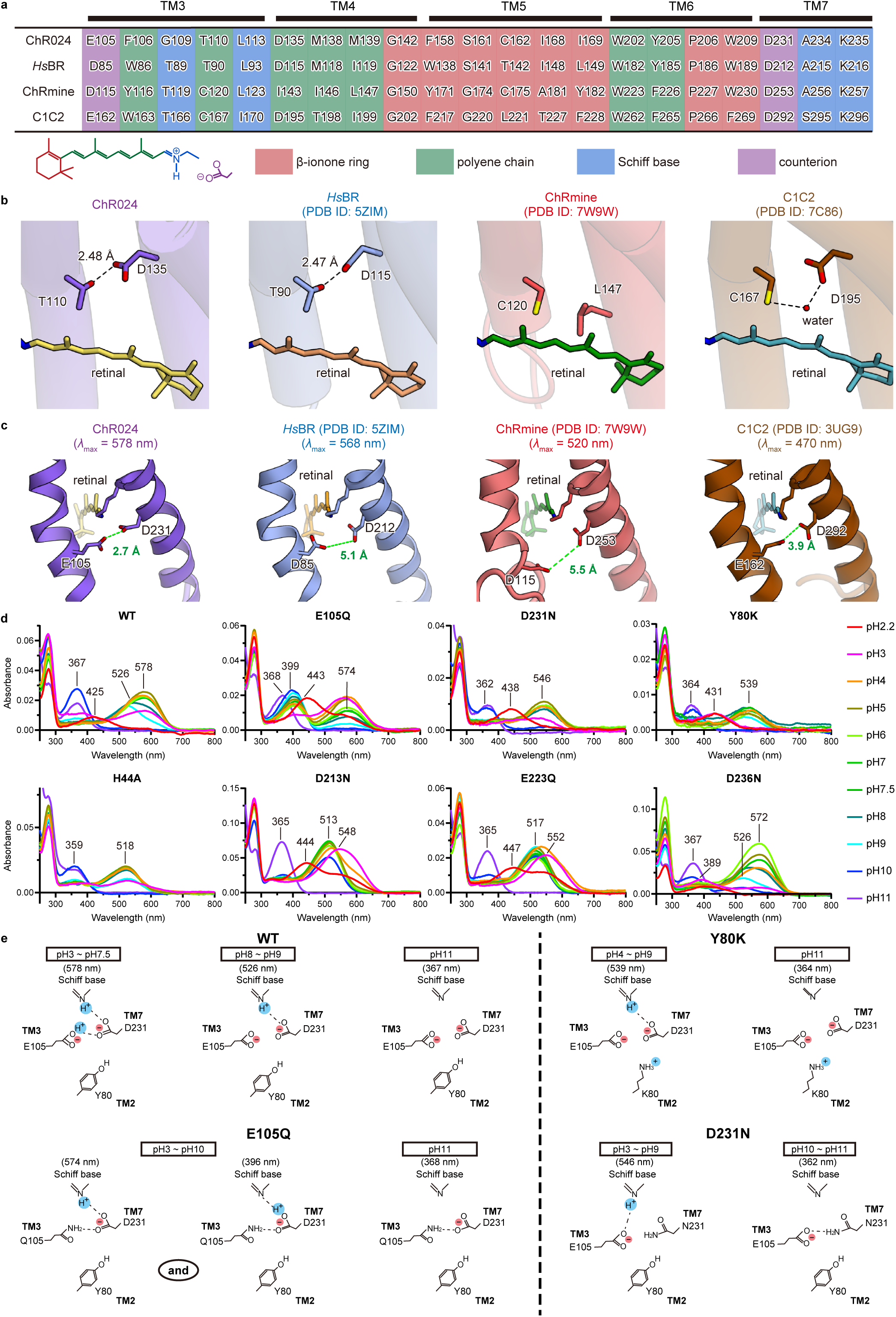
| Structural and spectroscopic analyses related to color tuning and channel function. **a,** Sequence alignment of residues in the retinal-binding pocket of ChR024, *Hs*BR, ChRmine, and C1C2. Each residue in the retinal binding pocket is color-coded according to the position of retinal: counterion residues are shown in purple, residues around the β-ionone ring in red, those around the polyene chain in green, and those around the Schiff base in blue. **b,** Geometry of counterions in TM3 and TM7. Green dashed lines indicate the shortest distance between the counterions. **c,** Geometry of residues at positions corresponding to the so called “DC gate” in canonical channelrhodopsins. The model with PDB ID 7C86^37^ was used to depict the water molecule located between the DC gate residues instead of 3UG9.^82^ Note that in the density map of 3UG9, this water molecule was not modeled, but weak water-like density was present at the corresponding position, supporting the validity of this substitution. **d,** pH-dependent absorption spectra of ChR024 WT and the mutants E105Q, D231N, Y80K, H44A, D213N, E223Q, and D236N, measured from pH 2.2 to 11.0. **e,** Schematic models of the ionic state of the Schiff base, the counterions, and the p*K*_a_-regulating residues at position 80 in TM2 for ChR024 WT and the mutants Y80K, E105Q, and D231N under different pH conditions.

**Supplementary Fig. 4.**
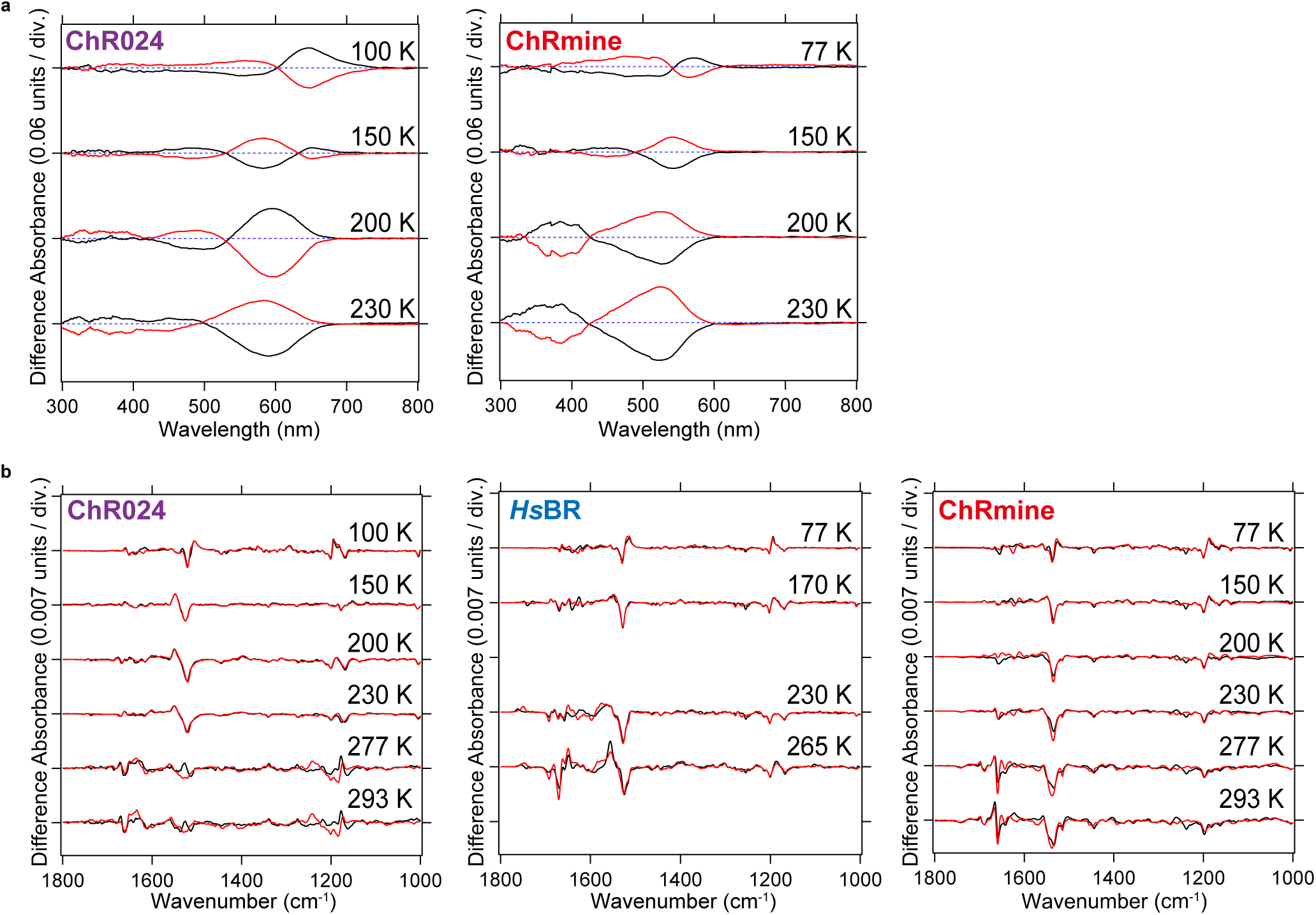
| Spectroscopic characterization of ChR024, *Hs*BR, and ChRmine. **a,** Light-induced low-temperature difference UV-visible spectra of ChR024 (left) obtained at 100, 150, 200, and 230 K. Black curves represent difference spectra recorded after illumination at 570 nm for 2 min (100 K) or >550 nm for 2 min (150, 200, 230 K). Red curves represent reversion from intermediates after illumination at >610 nm for 2 min (100 K) or 480 nm for 2 min (150, 200, 230 K). Light-induced low-temperature difference UV-visible spectra of ChRmine (right) obtained at 77, 150, 200, 230 K. Black curves represent difference spectra after illumination at 520 nm for 2 min (77 K) or >500 nm for 2 min (150, 200, 230 K). Red curves represent reversion from intermediates after illumination at >550 nm for 2 min (100 K), 460 nm for 2 min (150 K), or 400 nm for 2 min (200, 230 K). **b,** Light-induced low-temperature difference FTIR spectra of ChR024 (left), *Hs*BR (middle), and ChRmine (right) in H_2_O (black) and D_2_O (red), in the 1800–1000 cm^-1^ region. The FTIR spectra of *Hs*BR were reproduced from Ref 44–47.

