## Supplementary Notes for "Structural basis for color tuning and passive ion conductance in red-shifted pump-fold channelrhodopsin ChR024"

### Protonation States of H44 and D236

Our calculations initially assigned the protonated and deprotonated states to H44 and D236, respectively. This assignment enabled us to extract H44 and D236, together with K154, R212, D213, and E223, as key residues responsible for color tuning in ChR024 (Fig. 3b). However, under this assumption, the calculations predicted that the H44A and D236N mutations would increase and decrease the p$K_a$ of E105 by +5.6 and –2.5, respectively. These predictions are inconsistent with the experimentally measured $\lambda_{max}$ values, which indicate that E105 is deprotonated in H44A and protonated in D236N (Fig. 3c and Supplementary Fig. 3e). We therefore attribute this discrepancy to the limited accuracy of the protonation state model employed in our calculations.

In general, protonation states can be evaluated by representing the protons of the titrating side-chains explicitly or implicitly.[1] In the present work, we modeled the titrating protons of Asp and Glu side-chains implicitly by rearranging the partial charges on the heavy (non-hydrogen) atoms, as is commonly done in a simple electrostatic continuum model. This model does not fully account for stabilization by specific hydrogen bonds involving the titrating protons of Asp and Glu side-chains. Structural inspection suggests that the $N_\varepsilon$ atom of H44 is located 2.7 Å from deprotonated D231, supporting its protonated state. In contrast, the $N_\delta$ atom of H44 lies 2.8 Å from D236, giving rise to two plausible scenarios: the salt-bridge state [H44–$N_\delta H^+ \cdots O_{\delta 1}$–D236$^-$] and the neutral–neutral state [H44–$N_\delta \cdots HO_{\delta 1}$–D236]. Because the implicit model cannot capture stabilization arising from the hydrogen bond in the neutral–neutral configuration, the free energy of this state was likely overestimated. As a result, the calculations favored the salt-bridge state [H44–$N_\delta H^+ \cdots O_{\delta 1}$–D236$^-$].


1. Simonson, T., Carlsson, J. & Case, D. A. Proton Binding to Proteins: p$K_a$ Calculations with Explicit and Implicit Solvent Models. *J. Am. Chem. Soc.* **126**, 4167–4180 (2004).
