## Supplementary Table 1 for "Structural basis for color tuning and passive ion conductance in red-shifted pump-fold channelrhodopsin ChR024"

### Cryo-EM data collection, refinement and validation statistics

| | ChR024 in DDM micelles (EMD-XXXXX) | ChR024 in MSP1E3D1 nanodiscs (EMD-XXXXX) (PDB XXXX) |
|---|---|---|
| **Data collection and processing** | | |
| Magnification | 105,000 | 105,000 |
| Voltage (kV) | 300 | 300 |
| Electron exposure (e–/Å$^2$) | 46 | 45.2 |
| Defocus range (μm) | -0.8 ~ -1.6 | -0.8 ~ -1.6 |
| Pixel size (Å) | 0.83 | 0.83 |
| Symmetry imposed | C3 | C3 |
| Final particle images (no.) | 342,264 | 415,817 |
| Map resolution (Å) | 3.22 | 2.44 |
| FSC threshold | 0.143 | 0.143 |
| **Refinement** | | |
| Initial model used (PDB code) | | AF2-generated model |
| Map sharpening *B* factor (Å$^2$) | | -92.9 |
| Model composition (monomer) | | |
| Non-hydrogen atoms | | 1863 |
| Protein residues | | 226 |
| Retinal | | 20 |
| Water | | 16 |
| *B* factors (Å$^2$) | | |
| Protein | | 74.9 |
| Retinal | | 52.0 |
| Water | | 74.6 |
| R.m.s. deviations | | |
| Bond lengths (Å) | | 0.0062 |
| Bond angles (°) | | 1.476 |
| Validation | | |
| MolProbity score | | 1.63 |
| Clashscore | | 3.51 |
| Poor rotamers (%) | | 2.51 |
| Ramachandran plot | | |
| Favored (%) | | 96.88 |
| Allowed (%) | | 3.12 |
| Disallowed (%) | | 0 |
